# Protonation-Dependent Sequencing of 5-Formylcytidine in RNA

**DOI:** 10.1101/2021.11.23.469744

**Authors:** Courtney N. Link, Supuni Thalalla Gamage, Diamond A. Gallimore, Robert Kopajtich, Christine N. Evans, Samantha R. Nance, Stephen D. Fox, Thorkell Andresson, Raj Chari, Joseph Ivanic, Holger Prokisch, Jordan L. Meier

## Abstract

Chemical modification of cytidine in non-coding RNAs plays a key role in regulating translation and disease. However, the distribution and dynamics of many of these modifications remains unknown due to a lack of sensitive site-specific sequencing technologies. Here we report a protonation-dependent sequencing reaction for detection of 5-formylcytidine (5fC) and 5-carboxycytidine (5caC) in RNA. First, we evaluate how protonation combined with electron-withdrawing substituents alters the molecular orbital energies and reduction of modified cytidine nucleosides, highlighting 5fC and 5caC as reactive species. Next, we apply this reaction to detect these modifications in synthetic oligonucleotides as well as endogenous human tRNA. Finally, we demonstrate the utility of our method to characterize a patient-derived model of 5fC-deficiency, where it enables facile monitoring of both pathogenic loss and exogenous rescue of NSUN3-dependent 5fC within the wobble base of human mitochondrial tRNA^Met^. These studies showcase the ability of protonation to enhance the reactivity and sensitive detection of 5fC in RNA, and provide a molecular foundation for applying optimized sequencing reactions to better understand the role of oxidized RNA cytidine nucleobases in disease.

## Introduction

Chemical modification of coding and non-coding nucleic acids plays a crucial role in gene regulation and cellular homeostasis.^1,2^ In human RNA, the methylated nucleobase 5-methylcytidine (5mC) occurs in transfer RNA (tRNA), ribosomal RNA (rRNA), and messenger RNA (mRNA). Deposition of this modification into RNA is regulated by DNMT2 and the NSUN methyltransferase family and has been linked to protein translation, stress response, and differentiation.^3^ In addition to its intrinsic roles, 5mC can also serve as an intermediate in the formation of RNA 5-hydroxymethylcytidine (5hmC) and 5-formylcytidine (5fC) residues.^4,5^ Within DNA, 5hmC, 5fC, and the more highly oxidized 5-carboxycytidine (5caC) are well-known for their role in epigenetic regulation of gene expression. However, within RNA, the distribution and function of these oxidized cytidine derivatives is only beginning to emerge.

A key technical challenge to studying RNA modifications is the development of methods to reliably assess their location and penetrance at single nucleotide resolution.^6,7^ To date, the most widely applied protocols to study 5mC and its oxidized variants have employed bisulfite sequencing. This approach is based on the known resistance of 5mC and 5hmC to bisulfite-catalyzed deamination, a phenomenon that can be extended to 5fC and 5caC using chemical derivatization.^8–10^ One limitation of bisulfite-sequencing is that it creates signals at unmodified, rather than modified, cytidine nucleobases. This presents a challenge to the analysis of RNA, as loss of signal could either reflect the presence of an oxidized m5C or, alternatively, a highly structured region recalcitrant to bisulfite reactivity.^8,11^ Alternative protocols have emerged to that enable sensitive profiling of 5mC, 5hmC, and 5fC in DNA, but rely on custom synthetic reagents or enzymes that are not active on RNA.^12,13^ There remains an unmet need for sensitive methods to produce a quantitative signal specifically at oxidized cytidine derivatives in RNA.

Considering this challenge, we were inspired by the recent development of an optimized reaction for nucleotide resolution mapping of N4-acetylcytidine (ac4C) in RNA (Figure 1a).^14^ This reaction exploits the fact that N4-acetylation withdraws electron-density from the cytidine ring, rendering it susceptible to reduction by hydride donors. Optimization found the reaction can be further accelerated ∼10-fold by performing the reduction under strongly acidic conditions (pH 1).^14^ This protonates N3 and is believed to render the vinylic 5,6-double bond of cytidine hyperreactive. Reverse transcription (RT) using a processive polymerase (TGIRT) causes a non-cognate A to be introduced across from reduced ac4C during complementary DNA (cDNA) synthesis, which is read out as a C to T misincorporation at ac4C sites by sequencing.^15^ Interestingly, during the initial application of this reaction to the transcriptome-wide analysis of human RNA, we noted a minor site of cross-reactivity a known site of 5fC in mitochondrial tRNA^Met^.^16^ Hypothesizing this could be due to the electron-withdrawing properties of the 5-formyl group, we set out to explore the scope of protonation-dependent sequencing reactions for detection of cytidine modifications in RNA (Figure 1a).

**Figure 1.**
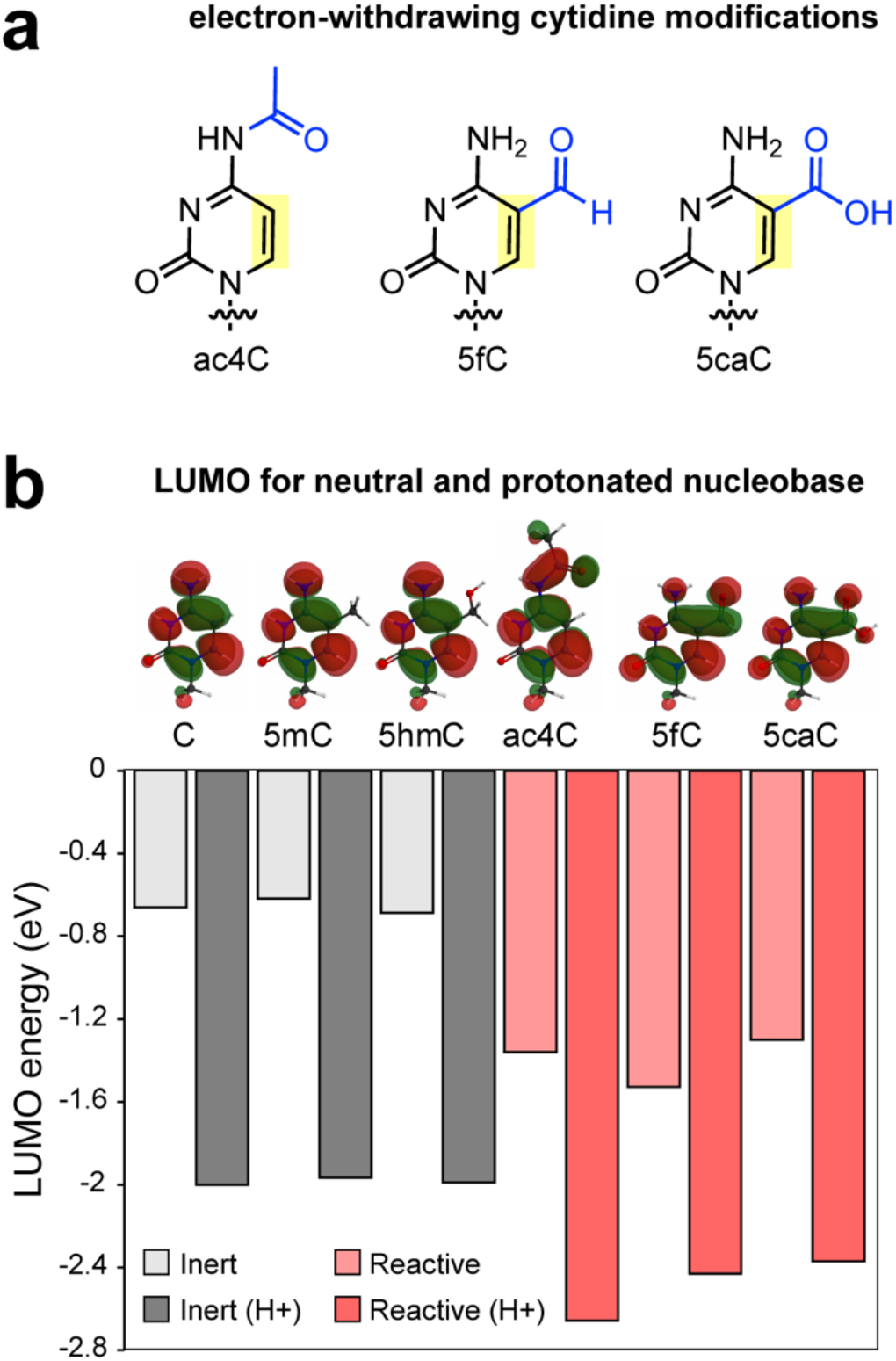
(a) Structure of modified cytidines containing electron-withdrawing groups. (b) Lowest unoccupied molecular orbital (LUMO) energies for modified cytidines examined in this study. Inert and reactive indicate nucleosides that were either inert or reactive in UV reactivity assay (Figure 2). Illustrated LUMOs are for protonated (H+) state. Additional computational chemistry data and procedures are provided in the Supporting Information.

## Results

To understand their potential to react with hydride donors, we first calculated molecular orbital energies for cytidine and its naturally-occurring modified analogues (Figure 1b, Figure S1). Consistent with prior reports, methylation (m5C) and hydroxymethylation (hm5C) had a minor effect on the lowest unoccupied molecular orbital (LUMO), which occurs across the 5,6-double bond and represents the site of hydride addition. Additional modifications such as 5-chlorocytidine (5ClC), N4-methylcytidine (m4C), N3-methylcytidine (m3C) also showed minor effects (Figure S1). In contrast, 5-formylation, 5-carboxylation (5caC), and N4-acetylation (ac4C) lowered the LUMO energy, which was further reduced by N3 protonation (Figure 1b). These calculations support the notion that electron-withdrawing modifications may alter the protonation-dependent reactivity of cytidine residues in RNA.

Next we experimentally assess the reactivity of modified cytidines using UV-spectrophotometry. This assay monitors the loss of absorbance caused by reduction and dearomatization of the cytidine ring in free nucleosides, allowing for facile reaction monitoring (Figure 2a).^15^ We focused our studies on 5mC, 5hmC, 5fC, and 5caC, with ac4C and cytidine serving as reactive and non-reactive control nucleobases, respectively (Figure 2b-c, Figure S2). In line with previous results, sodium cyanoborohydride (NaCNBH_3_) reduced ac4C in a pH-dependent manner over a time course of 2 h (Figure 2c, Figure S2). Under identical conditions the absorbance of cytidine was unaffected. Similarly, 5mC and 5hmC did not display evidence of NaCNBH_3_ reactivity at any pH. In contrast, more highly oxidized analogues demonstrated a strong reduction in absorbance upon treatment with hydride donors following the rank order 5fC > 5caC >> 5ClC. Deoxy-caC was also reduced, suggesting reactivity is independent of the nucleobase sugar (Figure S2). Furthermore, in each case this apparent reactivity was augmented by decreased (more acidic) pH. Previous studies have experimentally determined the pKa of protonated 5fC to be ∼1.5,^17^ and the greater reactivity of 5fC at pH 1 is consistent with the ability of N3 protonation to lower the barrier for hydride addition to the nucleobase. Hydride reduction of 5caC was observed to produce dihydrouridine, consistent with reduction, deamination, and decarboxylation (Figure 3a, Figure S3).^18^ In contrast, upon hydride reduction of 5fC dihydro-5-hydroxymethyluridine was the main uridine product formed, suggestive of reduction at both the 5,6 and 5-formyl positions and deamination (Figure S4). These results extend the scope of a pH-dependent reductive sequencing reaction to the 5fC and 5caC nucleobases.

**Figure 2.**
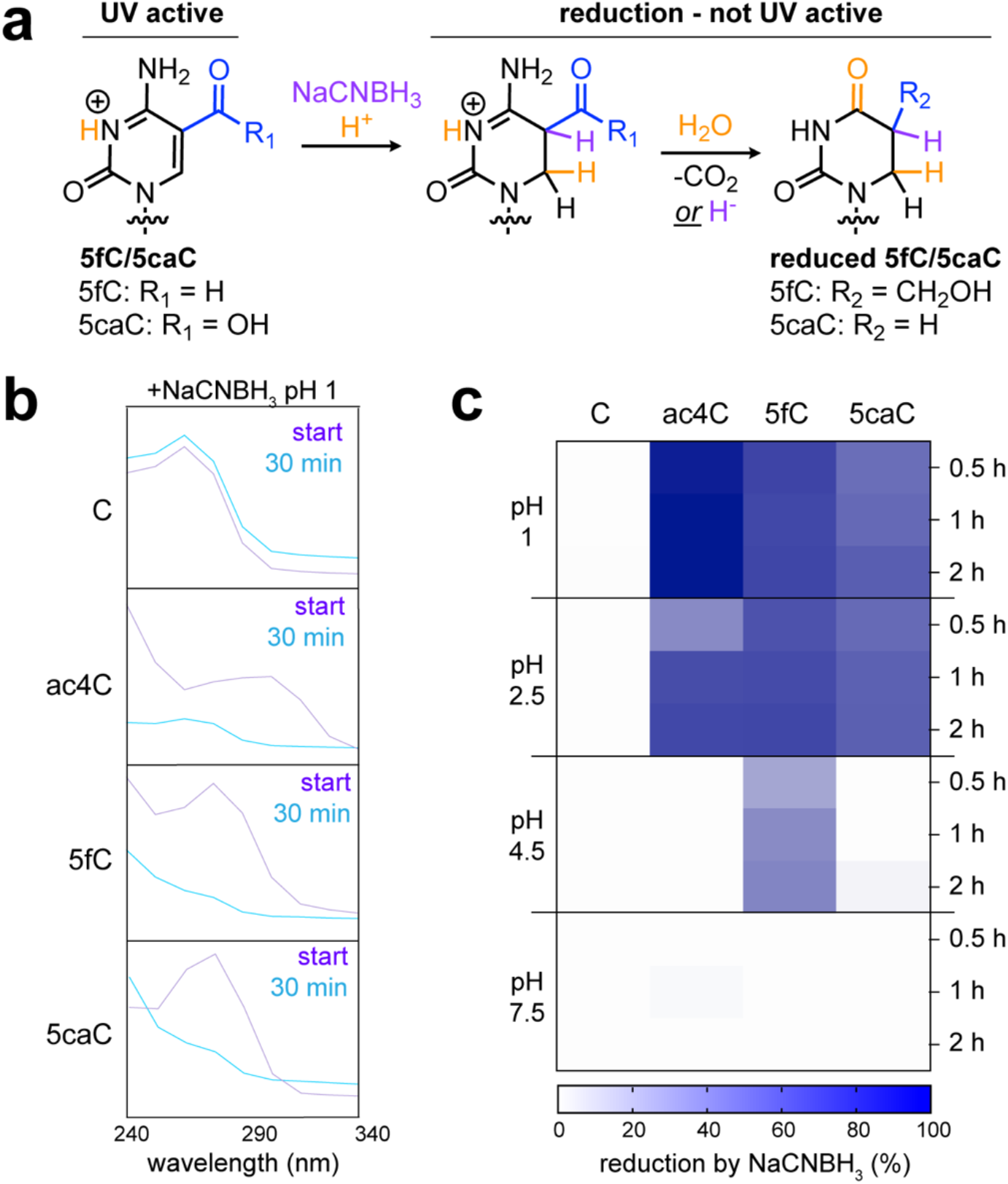
(a) Protonation-dependent reduction of 5fC and 5caC. This reaction can be followed by the conversion of UV absorbing starting materials to non-UV absorbing products. Mass spectrometry indicates 5fC and 5caC form dihydro-5-methyluridine and dihydrouridine, respectively (Figure S3-4). (b) Absorbance spectra for each nucleoside before and after incubation with NaCNBH_3_ (pH 1, 0.5 h). Graphs for additional time points and pH values are provided in Supporting Information Figure S2 (c) Heat map indicating percent of nucleoside consumed by NaCNBH_3_ as a function of pH and time.

**Figure 3.**
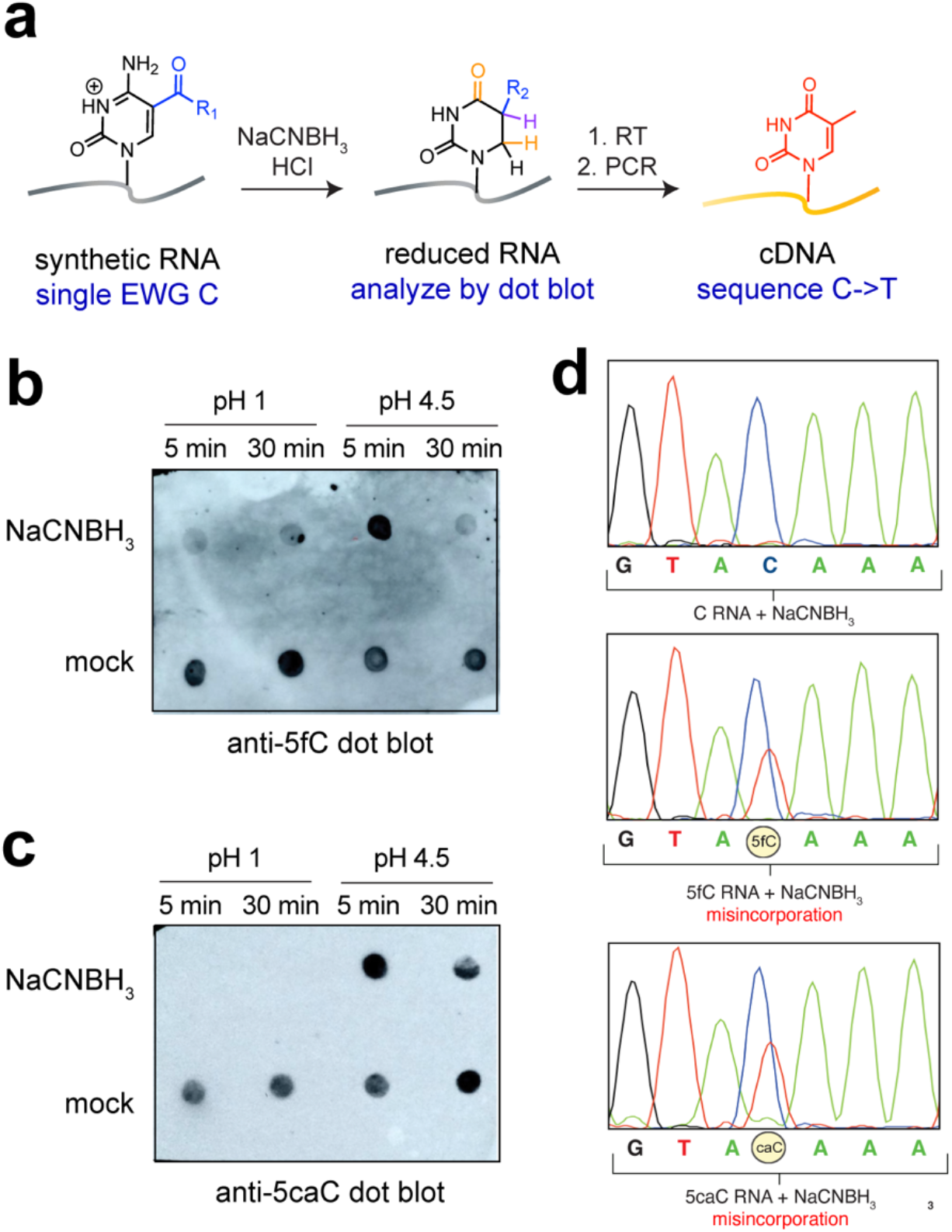
(a) Schematic for assessing reaction-based detection of 5caC and 5fC in RNA using protonation-assisted cyanoborohydride sequencing. (b) pH-dependent reaction of 5fC in RNA with NaCNBH_3_ as assessed by dot blot. (c) pH-dependent reaction of 5caC in RNA NaCNBH_3_as assessed by dot blot. (d) Misincorporation signals in RNAs containing C (top), 5fC (middle), and 5caC (bottom) following treatment with NaCNBH_3_ (pH 1), reverse transcription, PCR, and Sanger sequencing.

To effectively report on a modification’s presence, a sequencing reaction must alter the reverse transcription of RNA.^6^ Therefore, we next determined the pH-dependent reactivity of 5fC and 5caC in an RNA polynucleotide context. To enable these studies we first used in vitro transcription to produce a small panel of synthetic RNA substrates containing a single templated cytidine residue.^15^ (Figure 3a) Substituting modified cytidine triphosphates (e.g. 5fCTP, 5-caCTP) for CTP in the reaction enabled the preparation of a series of discretely modified synthetic RNAs. Using commercially-available 5fC and 5caC antibodies, we observed evidence for time-dependent reactivity of each nucleobase, with 5caC showing a greater dependence on acidic pH (Figure 3b-c). Galvanized by these studies, we assessed the ability of this reaction to cause RT-dependent misincorporation signals. For these studies RNA model substrates were treated with NaCNBH_3_ at the specified pH and reverse-transcribed using the processive TGIRT RT. Full length cDNA substrates were then PCR amplified, gel-extracted, and analyzed by Sanger sequencing. Focusing first on 5fC, we observed a borohydride-dependent mismatch at the 5fC site which was not observed in the cytidine, 5mC, or 5hmC substrates (Figure 3d, top and middle, Figure S5). Misincorporation at this site was partial (∼29%) and manifested as a C to U transversion, reflecting a misincorporation of adenosine into the antisense cDNA strand opposite the reduced 5fC (sense strand) during RT. A slightly higher (∼33%) misincorporation signal was observed for 5caC RNA (Figure 3d, bottom). This contrasts with the rank order reduction of the two nucleobases in our UV analysis, and may indicate differences in reactivity or relative ‘mutagenicity’ of the reduced nucleobases during RT.

Considering the potential for strong acid to catalyze RNA degradation, we next explored the interplay between 5fC conversion and hydrolysis. In choosing a milder reaction condition we were inspired by recent reports using 2-picoline borane reduction (pic-borane, pH 5.2) for the sequencing-based detection of 5fC and 5caC in DNA.^18^ Assessing the effect of the two reaction conditions on integrity of 28S and 18S rRNA in total RNA over 30 min, we observed pic-borane (pH 5.2) caused substantially less degradation than acidic NaCNBH_3_ (pH 1; Figure S6a). However, this was accompanied by reduced misincorporation at 5fC, despite heating and addition of 10-fold more reducing agent (Figure S6b). Extended treatment (3 h) by pic-borane at the milder pH led to similar levels of degradation as by NaCNBH_3_ (pH 1) (Figure S6a). The ability to detect 5fC despite fragmentation at acidic pH is consistent with our prior application of similarly acidic reaction conditions for ac4C sequencing. Overall these analyses define a kinetically-optimized chemistry for sensitive, RT-based detection of 5fC and 5caC in RNA.

Next, we sought to exploit this chemical reaction to detect oxidized electron-withdrawing cytidine modifications in endogenous RNA. Since RNA-associated 5caC has not yet been reported, we focused our efforts on 5fC. The human mitochondrial genome encodes a single copy of tRNA^Met^ which functions as both the initiator and elongator tRNA during mitochondrial protein synthesis (Figure 4a).^5,17^ Internal methionine incorporation requires the tRNA^Met^ CAU anticodon to recognize an AUA codon in the mRNA coding sequence, forming an unconventional wobble pair that is facilitated by 5fC. In addition to its unique coding properties, mitochondrial tRNA^Met^ is one of the only human tRNAs that does not contain any ‘hard stop’ base modifications that interfere with the canonical base pairing face of RNA (e.g. N7-methylguanosine or N1-methyladenine), suggesting its analysis may be particularly well-suited to our RT-dependent misincorporation approach.^19^ To test this, we subjected total RNA (∼5 ug) isolated from HEK-293T cells to parallel NaCNBH_3_ and control treatments reverse transcribed each sample using a primer specific to mitochondrial tRNA^Met^. The cDNA produced was then purified, PCR amplified, and analyzed by sequencing. Using this approach, we observed a partial (∼27%) misincorporation signal at position 34 of tRNA^Met^, corresponding to the wobble position (Figure 4b). Signal correlated with 5fC protonation, as less acidic conditions (NaCNBH_3_ pH 4.5, 30 min) produced a weaker signal (∼3%) (Figure S7). No signal was observed at neighboring cytidine residues or in the non-reduced control sample. These studies establish the feasibility of using a protonation-assisted sequencing reaction for the sensitive detection of endogenous electron-withdrawing cytidine modifications in RNA.

**Figure 4.**
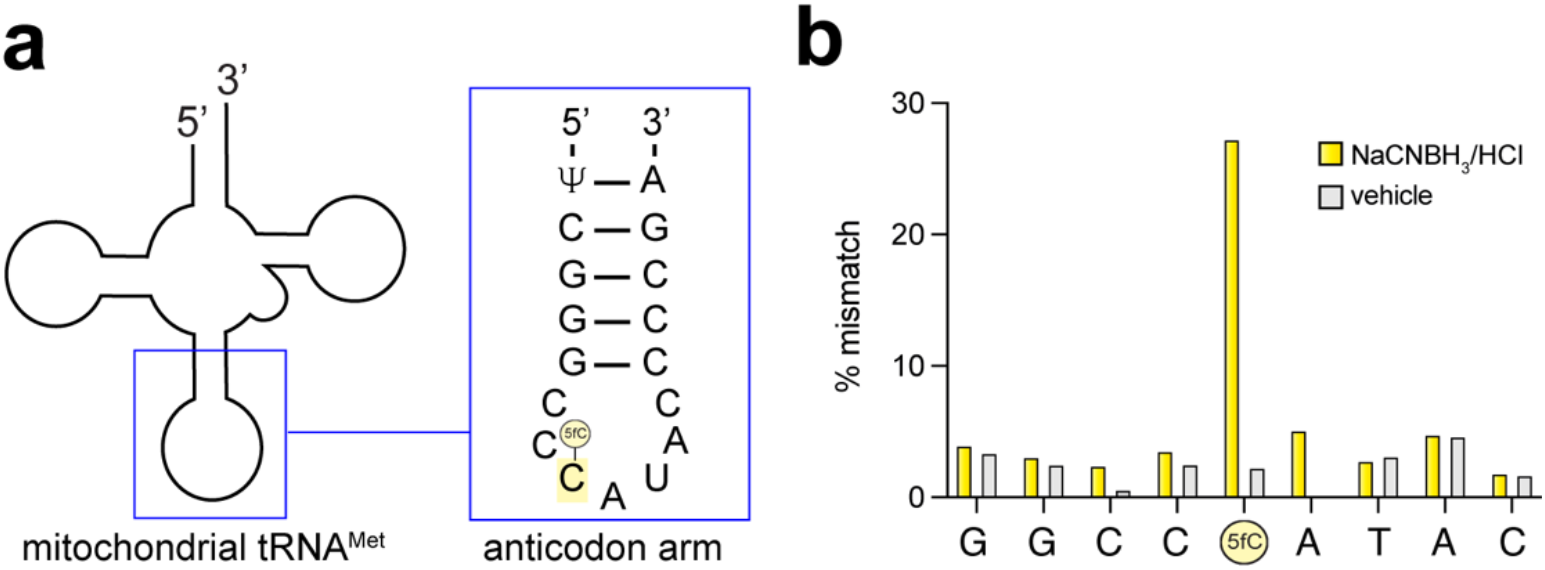
(a) Schematic of human mitochondrial tRNA^Met^ and sequence of anticodon arm. (b) Sequencing-based detection of 5fC at C34 of human mitochondrial tRNA^Met^ in endogenous RNA isolated from HEK-293T cells. Values represent averages of three independent technical replicates.

Mass spectrometry of RNA isolated from human placenta and HeLa cells have indicated that under basal conditions position 34 of mitochondrial tRNA^Met^ is modified with 5fC to a near uniform extent.^19^ Prior studies have found this owes to the dual activities of NSUN3 and ALKBH1, which methylate and oxidize the wobble base, respectively (Figure 5a).^20–22^ Inactivating mutations in the methyltransferase NSUN3 disrupt this process and are associated with mitochondrial respiratory chain complex deficiency.^23^ To assess whether our method could be used to monitor pathophysiological changes in NSUN3 or ALKBH1 activity, we first assessed the quantitative properties of our sequencing reaction. Model RNA substrates were prepared harboring either cytidine or 5fC, mixed in different proportions (100:0, 80:20, 60:40, etc), and subjected to our modification sequencing protocol. A linear relationship between 5fC level and C to T transversion at 5fC site was observed (R^2^ = 0.98; Figure 5b). This suggests the ability of our method to quantitatively assess 5fC levels, with the caveat that the absolute magnitude of misincorporation may be dependent on polynucleotide context. Galvanized by these results, we next applied our method to analyze RNA isolated from fibroblasts derived from patients harboring pathological NSUN3 mutations.^23^ Consistent with prior observations, we observed a complete loss of misincorporation signal at position 34 of tRNA^Met^ in RNA isolated from NSUN3 mutant patients as compared to wild-type (Figure 5c, Figure S8). Lentiviral re-introduction of NSUN3 into the mutant cell line rescued 5fC, albeit partially (Figure 5c, Figure S8). Of note, previous characterization of these models made use of bisulfite-sequencing based methods,^23^ and to our knowledge our study represents the first direct and quantitative assessment of 5fC in this pathophysiological setting.

**Figure 5.**
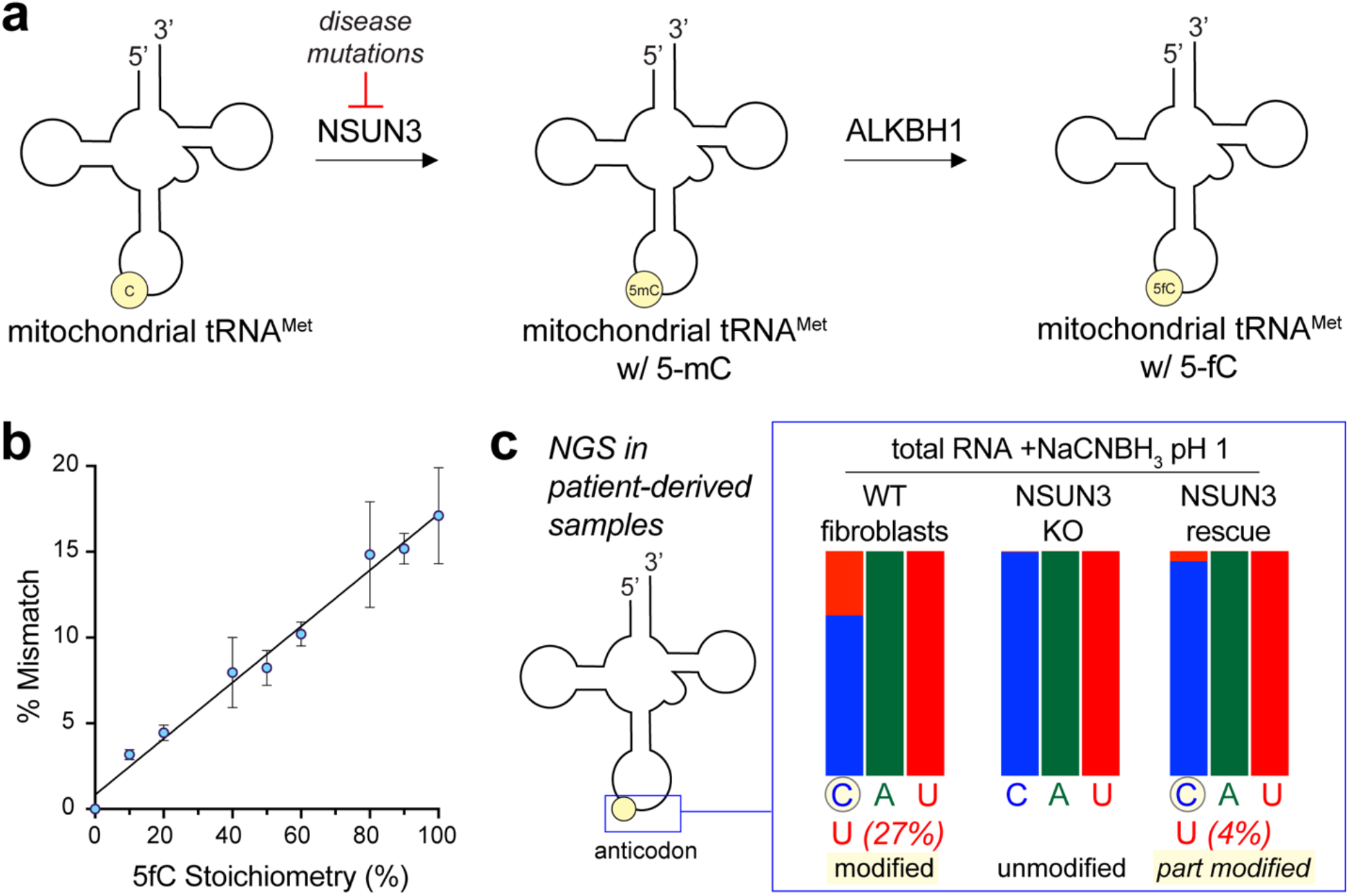
(a) Enzymes involved in biogenesis of 5fC at C34 mitochondrial tRNA^Met^. NSUN3 can be inactivated by pathogenic loss-of-function mutations. (b) Quantitative nature of 5fC sequencing reaction. Misincorporation was assessed following NaCNBH_3_ treatment, RT, and PCR of model RNAs containing defined stoichiometries of 5fC (0-100%) using Sanger sequencing. Values represent averages of two independent replicates. (c) Protonation-dependent 5fC sequencing can detect NSUN3 loss-of-function and rescue. RNA was isolated from wild-type human dermal fibroblasts (left), or fibroblasts derived from a patient harboring loss-of-function mutations in NSUN3 (middle), as well as a loss-of-function cell line in which NSUN3 had been re-introduced. NaCNBH_3_ treatment, RT, PCR, and targeted next-generation sequencing of the mitochondrial tRNA^Met^ amplicon differentiates the anticodon loop wild-type (left) and rescue (right) from NSUN3 knockout cells (middle). A “U” signal in the left-most nucleobase is indicative of 5fC’s presence. In all cases values are indicative of >10,000 reads at the specified positions.

## Discussion

Here we report the development of a protonation-assisted sequencing reaction to assess 5fC and 5caC in RNA. Molecular orbital analysis revealed N3 protonation lowers the LUMO of modified cytidines, which correlates with increased susceptibility to hydride reduction in model nucleosides and RNA substrates. Formation of the reduced nucleobases results in a misincorporation during reverse transcription, which can be detected via PCR amplification and sequencing. Acidic pH was found to be key to the kinetic optimization of this method, whose utility we demonstrate by monitoring an endogenous site of 5fC in both basal and pathophysiological settings. Our approach is differentiated from prior bisulfite-based methods to detect 5fC and 5caC by its ability to produce a signal specifically at the site of modification, rather than at unmodified cytidines. Another major advantage of the method is the signal enhancement afforded by PCR amplification, which allows quantitative analysis of 5fC in mitochondrial tRNA^Met^ using only micrograms of total RNA, as opposed to the milligrams of RNA or kilograms of tissue starting material required for mass spectrometry.^19,24^ The activation of ac4C, 5fC, and 5caC to nucleophilic attack by protonation has conceptual parallels in studies of small molecules whose electrophilicity is conditionally regulated by protonation or methylation,^25–27^ and it is possible that further exploration of this phenomenon may enable additional mapping applications. Relatedly, a key aspect of our method is the use of hydride donors that have a measurable half-life at low pH. A caveat to our method is that under our described protocol conversion of 5fC and 5caC was only partial, even at stoichiometrically-modified sites. In the future screening additional hydride donors, non-hydride nucleophiles, and reactant concentrations may yield more fully optimized reaction conditions. While our method produces signals directly at sites of electron-withdrawing cytidine modifications, it does not intrinsically differentiate ac4C, 5fC, and 5caC. Achieving this will require interfacing the current sequencing chemistry with additional control reactions, including hydrolytic lability of ac4C, imine formation with 5fC, and carbodiimide reactivity of 5caC.^9,10,15^

Finally, we note some future applications and areas for exploration. While we have demonstrated the ability of our sequencing method to monitor physiologically-relevant changes in 5fC, whether this modification is dynamically regulated by environmental stimuli or pathophysiological NSUN3, ALKBH1, or TET expression remains unknown.^28–30^ The sensitive and specific nature of our sequencing chemistry should greatly enable such studies. In addition, recent studies have suggested the potential for 5fC to occur in mRNA. We have previously applied reduction-based chemistry to identify RNA modification sites in sites eukaryotes and archaea,^16^ and we anticipate the current method should be well-suited to explore the distribution and prevalence of such modifications should they exist. Finally, our studies raise the potential of applying protonation-enhanced reduction chemistry to optimize protocols designed to profile 5mC and 5hmC in DNA, which previous studies have found can be detected by pic-borane sequencing following chemical or enzymatic oxidation.^18^ In model studies we have found reduction using NaCNBH_3_ (pH 1) can generate misincorporations at 5caC sites in a DNA substrate (Figure S9). Based on this, we anticipate our chemical insights should be readily integrable with existing protocols to oxidize and profile 5mC/5hmC, and potentially contribute to new methods for analysis of these marks in both DNA and RNA. Such studies are ongoing and will be reported in due course.

## Supporting Information

Supplemental figures, materials and methods are available online.

## Acknowledgements

The authors thank Prof. Schraga Schwartz (Weizmann Institute) for helpful discussions. This project has been funded in whole or in part with Federal funds from the National Cancer Institute, National Institutes of Health, under Contract No. HHSN261200800001E. This work was supported by the Intramural Research Program of the NIH, National Cancer Institute, Center for Cancer Research (ZIA-BC011488).

## References

(1) Cavalli, G.; Heard, E. Advances in Epigenetics Link Genetics to the Environment and Disease. Nature 2019, 571 (7766), 489–499.

(2) Frye, M.; Harada, B. T.; Behm, M.; He, C. RNA Modifications Modulate Gene Expression during Development. Science 2018, 361 (6409), 1346–1349.

(3) Popis, M. C.; Blanco, S.; Frye, M. Posttranscriptional Methylation of Transfer and Ribosomal RNA in Stress Response Pathways, Cell Differentiation, and Cancer. Curr. Opin. Oncol. 2016, 28 (1), 65–71.

(4) Fu, L.; Guerrero, C. R.; Zhong, N.; Amato, N. J.; Liu, Y.; Liu, S.; Cai, Q.; Ji, D.; Jin, S.-G.; Niedernhofer, L. J.; Pfeifer, G. P.; Xu, G.-L.; Wang, Y. Tet-Mediated Formation of 5-Hydroxymethylcytosine in RNA. J. Am. Chem. Soc. 2014, 136 (33), 11582–11585.

(5) Cantara, W. A.; Murphy, F. V., 4th; Demirci, H.; Agris, P. F. Expanded Use of Sense Codons Is Regulated by Modified Cytidines in tRNA. Proc. Natl. Acad. Sci. U. S. A. 2013, 110 (27), 10964–10969.

(6) Bartee, D.; Gamage, S. T.; Link, C. N.; Meier, J. L. Arrow Pushing in RNA Modification Sequencing. Chemical Society Reviews. 2021, pp 9482–9502. https://doi.org/10.1039/d1cs00214g.

(7) Wiener, D.; Schwartz, S. The Epitranscriptome beyond mA. Nat. Rev. Genet. 2021, 22 (2), 119–131.

(8) Khoddami, V.; Yerra, A.; Mosbruger, T. L.; Fleming, A. M.; Burrows, C. J.; Cairns, B. R. Transcriptome-Wide Profiling of Multiple RNA Modifications Simultaneously at Single-Base Resolution. Proc. Natl. Acad. Sci. U. S. A. 2019, 116 (14), 6784–6789.

(9) Booth, M. J.; Marsico, G.; Bachman, M.; Beraldi, D.; Balasubramanian, S. Quantitative Sequencing of 5-Formylcytosine in DNA at Single-Base Resolution. Nat. Chem. 2014, 6 (5), 435–440.

(10) Lu, X.; Song, C.-X.; Szulwach, K.; Wang, Z.; Weidenbacher, P.; Jin, P.; He, C. Chemical Modification-Assisted Bisulfite Sequencing (CAB-Seq) for 5-Carboxylcytosine Detection in DNA. J. Am. Chem. Soc. 2013, 135 (25), 9315–9317.

(11) Zhang, Z.; Chen, T.; Chen, H.-X.; Xie, Y.-Y.; Chen, L.-Q.; Zhao, Y.-L.; Liu, B.-D.; Jin, L.; Zhang, W.; Liu, C.; Ma, D.-Z.; Chai, G.-S.; Zhang, Y.; Zhao, W.-S.; Ng, W. H.; Chen, J.; Jia, G.; Yang, J.; Luo, G.-Z. Systematic Calibration of Epitranscriptomic Maps Using a Synthetic Modification-Free RNA Library. Nat. Methods 2021, 18 (10), 1213–1222.

(12) Schutsky, E. K.; DeNizio, J. E.; Hu, P.; Liu, M. Y.; Nabel, C. S.; Fabyanic, E. B.; Hwang, Y.; Bushman, F. D.; Wu, H.; Kohli, R. M. Nondestructive, Base-Resolution Sequencing of 5-Hydroxymethylcytosine Using a DNA Deaminase. Nat. Biotechnol. 2018. https://doi.org/10.1038/nbt.4204.

(13) Zeng, H.; He, B.; Xia, B.; Bai, D.; Lu, X.; Cai, J.; Chen, L.; Zhou, A.; Zhu, C.; Meng, H.; Gao, Y.; Guo, H.; He, C.; Dai, Q.; Yi, C. Bisulfite-Free, Nanoscale Analysis of 5-Hydroxymethylcytosine at Single Base Resolution. J. Am. Chem. Soc. 2018, 140 (41), 13190–13194.

(14) Thalalla Gamage, S.; Sas-Chen, A.; Schwartz, S.; Meier, J. L. Quantitative Nucleotide Resolution Profiling of RNA Cytidine Acetylation by ac4C-Seq. Nat. Protoc. 2021, 16 (4), 2286–2307.

(15) Thomas, J. M.; Briney, C. A.; Nance, K. D.; Lopez, J. E.; Thorpe, A. L.; Fox, S. D.; Bortolin-Cavaille, M.-L.; Sas-Chen, A.; Arango, D.; Oberdoerffer, S.; Cavaille, J.; Andresson, T.; Meier, J. L. A Chemical Signature for Cytidine Acetylation in RNA. J. Am. Chem. Soc. 2018, 140 (40), 12667–12670.

(16) Sas-Chen, A.; Thomas, J. M.; Matzov, D.; Taoka, M.; Nance, K. D.; Nir, R.; Bryson, K. M.; Shachar, R.; Liman, G. L. S.; Burkhart, B. W.; Gamage, S. T.; Nobe, Y.; Briney, C. A.; Levy, M. J.; Fuchs, R. T.; Robb, G. B.; Hartmann, J.; Sharma, S.; Lin, Q.; Florens, L.; Washburn, M. P.; Isobe, T.; Santangelo, T. J.; Shalev-Benami, M.; Meier, J. L.; Schwartz, S. Dynamic RNA Acetylation Revealed by Quantitative Cross-Evolutionary Mapping. Nature 2020, 583 (7817), 638–643.

(17) Lusic, H.; Gustilo, E. M.; Vendeix, F. A. P.; Kaiser, R.; Delaney, M. O.; Graham, W. D.; Moye, V. A.; Cantara, W. A.; Agris, P. F.; Deiters, A. Synthesis and Investigation of the 5-Formylcytidine Modified, Anticodon Stem and Loop of the Human Mitochondrial tRNAMet. Nucleic Acids Res. 2008, 36 (20), 6548–6557.

(18) Liu, Y.; Siejka-Zielińska, P.; Velikova, G.; Bi, Y.; Yuan, F.; Tomkova, M.; Bai, C.; Chen, L.; Schuster-Böckler, B.; Song, C.-X. Bisulfite-Free Direct Detection of 5-Methylcytosine and 5-Hydroxymethylcytosine at Base Resolution. Nat. Biotechnol. 2019, 37 (4), 424–429.

(19) Suzuki, T.; Yashiro, Y.; Kikuchi, I.; Ishigami, Y.; Saito, H.; Matsuzawa, I.; Okada, S.; Mito, M.; Iwasaki, S.; Ma, D.; Zhao, X.; Asano, K.; Lin, H.; Kirino, Y.; Sakaguchi, Y.; Suzuki, T. Complete Chemical Structures of Human Mitochondrial tRNAs. Nat. Commun. 2020, 11 (1), 4269.

(20) Haag, S.; Sloan, K. E.; Ranjan, N.; Warda, A. S.; Kretschmer, J.; Blessing, C.; Hübner, B.; Seikowski, J.; Dennerlein, S.; Rehling, P.; Rodnina, M. V.; Höbartner, C.; Bohnsack, M. T. NSUN3 and ABH1 Modify the Wobble Position of Mt-tRNAMet to Expand Codon Recognition in Mitochondrial Translation. EMBO J. 2016, 35 (19), 2104–2119.

(21) Kawarada, L.; Suzuki, T.; Ohira, T.; Hirata, S.; Miyauchi, K.; Suzuki, T. ALKBH1 Is an RNA Dioxygenase Responsible for Cytoplasmic and Mitochondrial tRNA Modifications. Nucleic Acids Res. 2017, 45 (12), 7401–7415.

(22) Nakano, S.; Suzuki, T.; Kawarada, L.; Iwata, H.; Asano, K.; Suzuki, T. NSUN3 Methylase Initiates 5-Formylcytidine Biogenesis in Human Mitochondrial tRNA(Met). Nat. Chem. Biol. 2016, 12 (7), 546–551.

(23) Van Haute, L.; Dietmann, S.; Kremer, L.; Hussain, S.; Pearce, S. F.; Powell, C. A.; Rorbach, J.; Lantaff, R.; Blanco, S.; Sauer, S.; Kotzaeridou, U.; Hoffmann, G. F.; Memari, Y.; Kolb-Kokocinski, A.; Durbin, R.; Mayr, J. A.; Frye, M.; Prokisch, H.; Minczuk, M. Deficient Methylation and Formylation of Mt-tRNA(Met) Wobble Cytosine in a Patient Carrying Mutations in NSUN3. Nat. Commun. 2016, 7, 12039.

(24) Suzuki, T.; Suzuki, T. Chaplet Column Chromatography: Isolation of a Large Set of Individual RNAs in a Single Step. Methods in Enzymology. 2007, pp 231–239. https://doi.org/10.1016/s0076-6879(07)25010-4.

(25) Kulkarni, R. A.; Bak, D. W.; Wei, D.; Bergholtz, S. E.; Briney, C. A.; Shrimp, J. H.; Alpsoy, A.; Thorpe, A. L.; Bavari, A. E.; Crooks, D. R.; Levy, M.; Florens, L.; Washburn, M. P.; Frizzell, N.; Dykhuizen, E. C.; Weerapana, E.; Linehan, W. M.; Meier, J. L. A Chemoproteomic Portrait of the Oncometabolite Fumarate. Nat. Chem. Biol. 2019, 15 (4), 391–400.

(26) Johnson, C. M.; Linsky, T. W.; Yoon, D.-W.; Person, M. D.; Fast, W. Discovery of Halopyridines as Quiescent Affinity Labels: Inactivation of Dimethylarginine Dimethylaminohydrolase. J. Am. Chem. Soc. 2011, 133 (5), 1553–1562.

(27) Schmidt, T. J.; Ak, M.; Mrowietz, U. Reactivity of Dimethyl Fumarate and Methylhydrogen Fumarate towards Glutathione and N-Acetyl-L-Cysteine--Preparation of S-Substituted Thiosuccinic Acid Esters. Bioorg. Med. Chem. 2007, 15 (1), 333–342.

(28) Paramasivam, A.; Meena, A. K.; Venkatapathi, C.; Pitceathly, R. D. S.; Thangaraj, K. Novel Biallelic NSUN3 Variants Cause Early-Onset Mitochondrial Encephalomyopathy and Seizures. Journal of Molecular Neuroscience. 2020, pp 1962–1965. https://doi.org/10.1007/s12031-020-01595-8.

(29) Zhang, Y.; Wang, C. Demethyltransferase AlkBH1 Substrate Diversity and Relationship to Human Diseases. Mol. Biol. Rep. 2021, 48 (5), 4747–4756.

(30) Yue, X.; Rao, A. TET Family Dioxygenases and the TET Activator Vitamin C in Immune Responses and Cancer. Blood. 2020, pp 1394–1401. https://doi.org/10.1182/blood.2019004158.

