## Supplementary Information for "Protonation-Dependent Sequencing of 5-Formylcytidine in RNA"

**Table of Contents for Supporting Information**

| | Page |
|---|---|
| Figure S1: LUMO energies for modified nucleobases. | S2 |
| Figure S2: Absorbance spectra for nucleosides before and after $NaCNBH_3$ | S3 |
| Figure S3: Analysis of 5-carboxycytidine products | S4 |
| Figure S4: Analysis of 5-formylcytidine products | S5 |
| Figure S5: 5mC and 5hmC are not reduced by $NaCNBH_3$ (pH 1) | S6 |
| Figure S6: Analysis of RNA degradation by sequencing conditions | S7 |
| Figure S7: pH dependence of tRNA 5fC conversion | S8 |
| Figure S8: Analysis of NSUN3 samples | S9 |
| Figure S9: Analysis of 5-carboxylcytidine in DNA | S10 |
| Experimental procedures and protocols | S11-15 |
| Table of DNA templates and primers | S16-17 |
| Raw computational chemistry data | S17-35 |
| References | S36 |

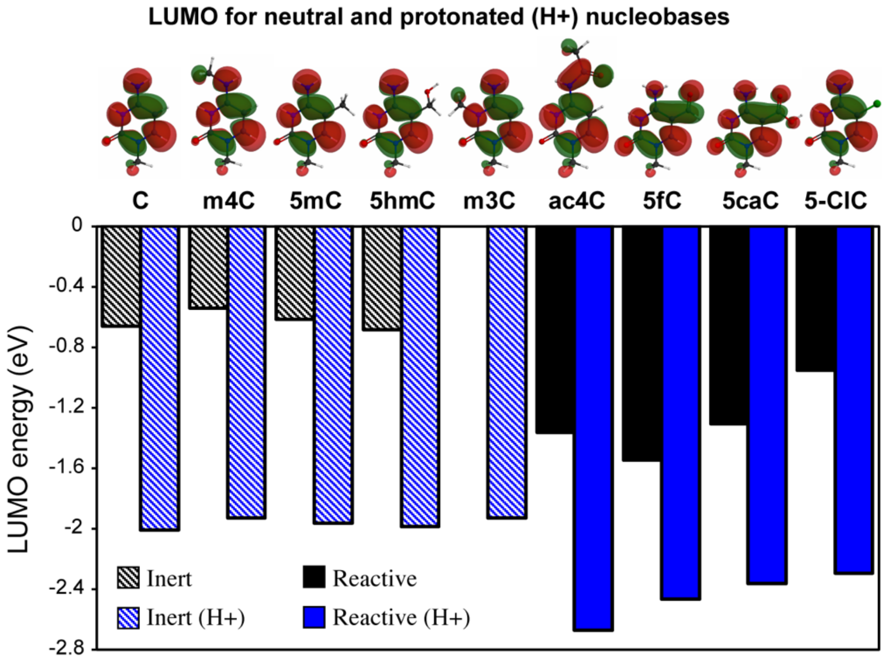


**Figure S1**. B3LYP-PCM($H_2O$) LUMOs shown for neutral and protonated (H+) systems. Inert and reactive indicate nucleosides that were either inert or reactive in UV reactivity assay (Figure 2), and illustrations are for protonated states. Computational chemistry procedures and data are provided in the Supporting Information (pages S11 and S17-35). C = cytidine, m4C = N4-methylcytidine, 5mC = 5-methylcytidine, 5hmC = 5-hydroxymethylcytidine, m3C = N3-methylcytidine, ac4C = N4-acetylcytidine, 5fC = 5-formylcytidine, 5caC = 5-carboxylcytidine, 5ClC = 5-chlorocytidine.

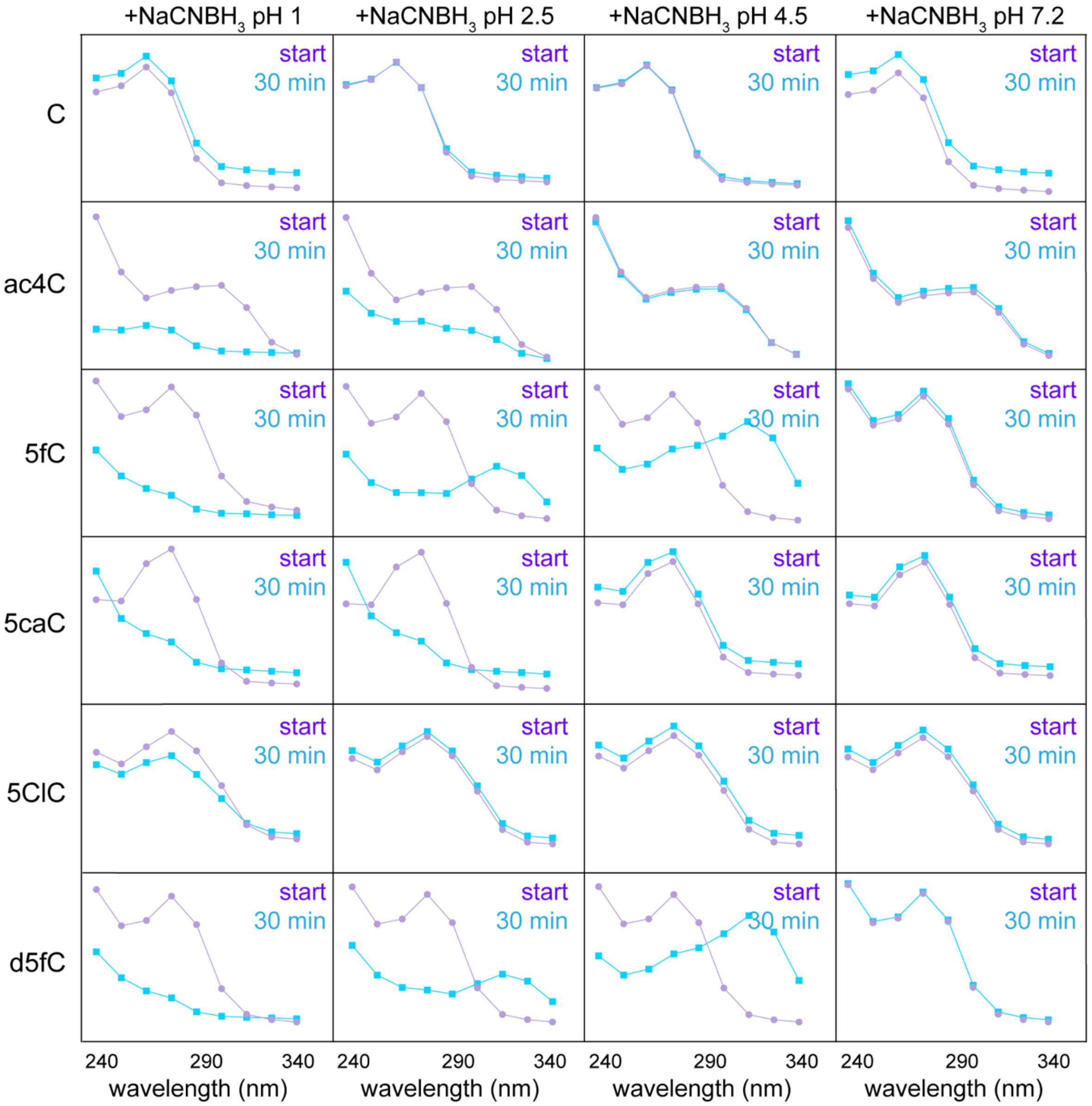


**Figure S2**. Absorbance spectra for each nucleoside before and after incubation with $NaCNBH_3$ as a function of pH. C = cytidine, ac4C = N4-acetylcytidine, 5fC = 5-formylcytidine, 5caC = 5-carboxylcytidine, 5ClC = 5-chlorocytidine, d5fC = deoxy-5-formylcytidine.

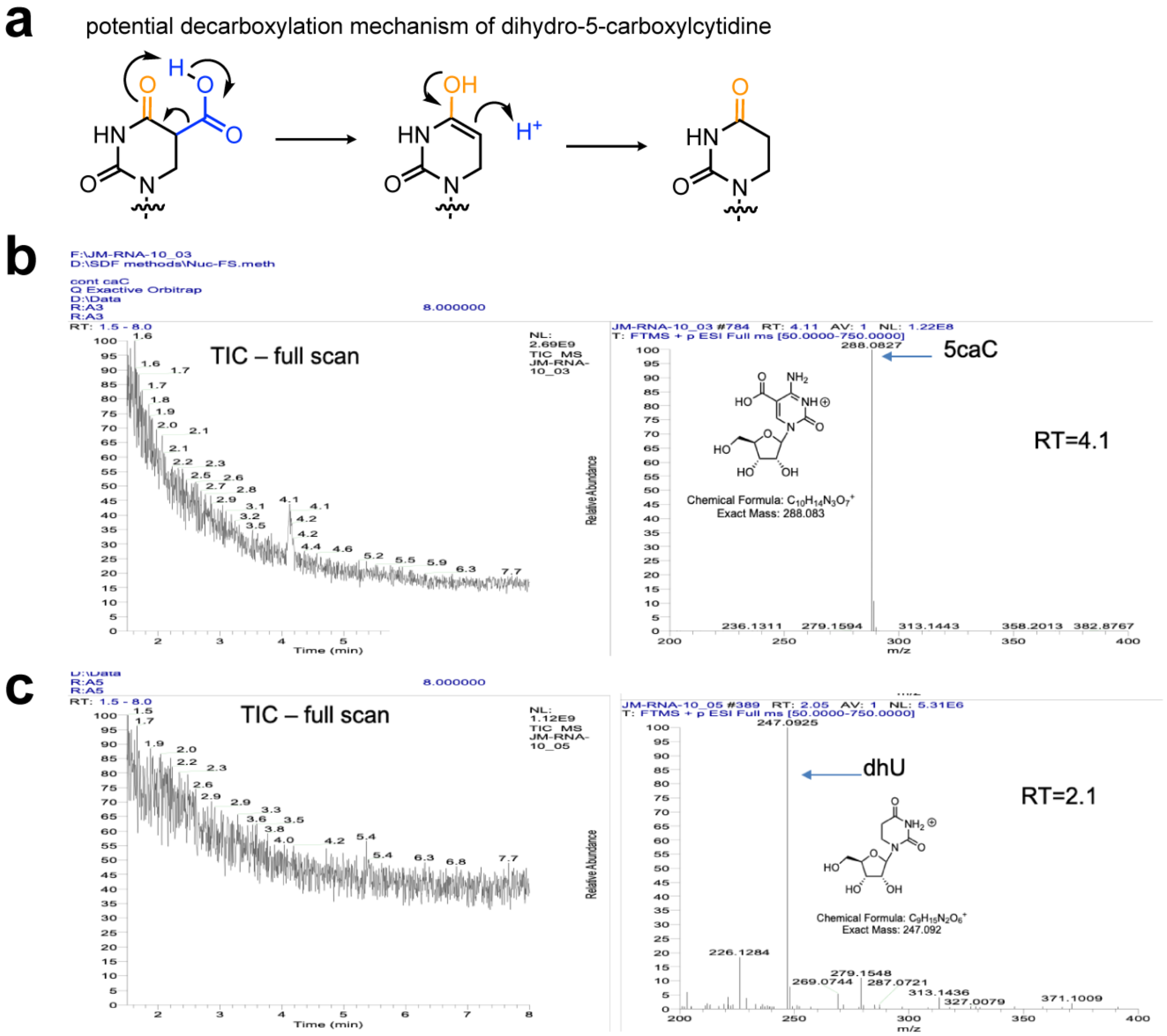


**Figure S3**. Product analysis of 5-carboxycytidine following $NaCNBH_3$ (pH 1) treatment. (a) Potential mechanism for conversion of 5-carboxy-dihydrouridine to dihydrouridine. This step would require prior reduction and deamination of caC. (b) LC-MS analysis of 5caC starting material. (c) LC-MS analysis indicating formation of dihydrouridine (dhU) from 5caC following $NaCNBH_3$ (pH 1) treatment.

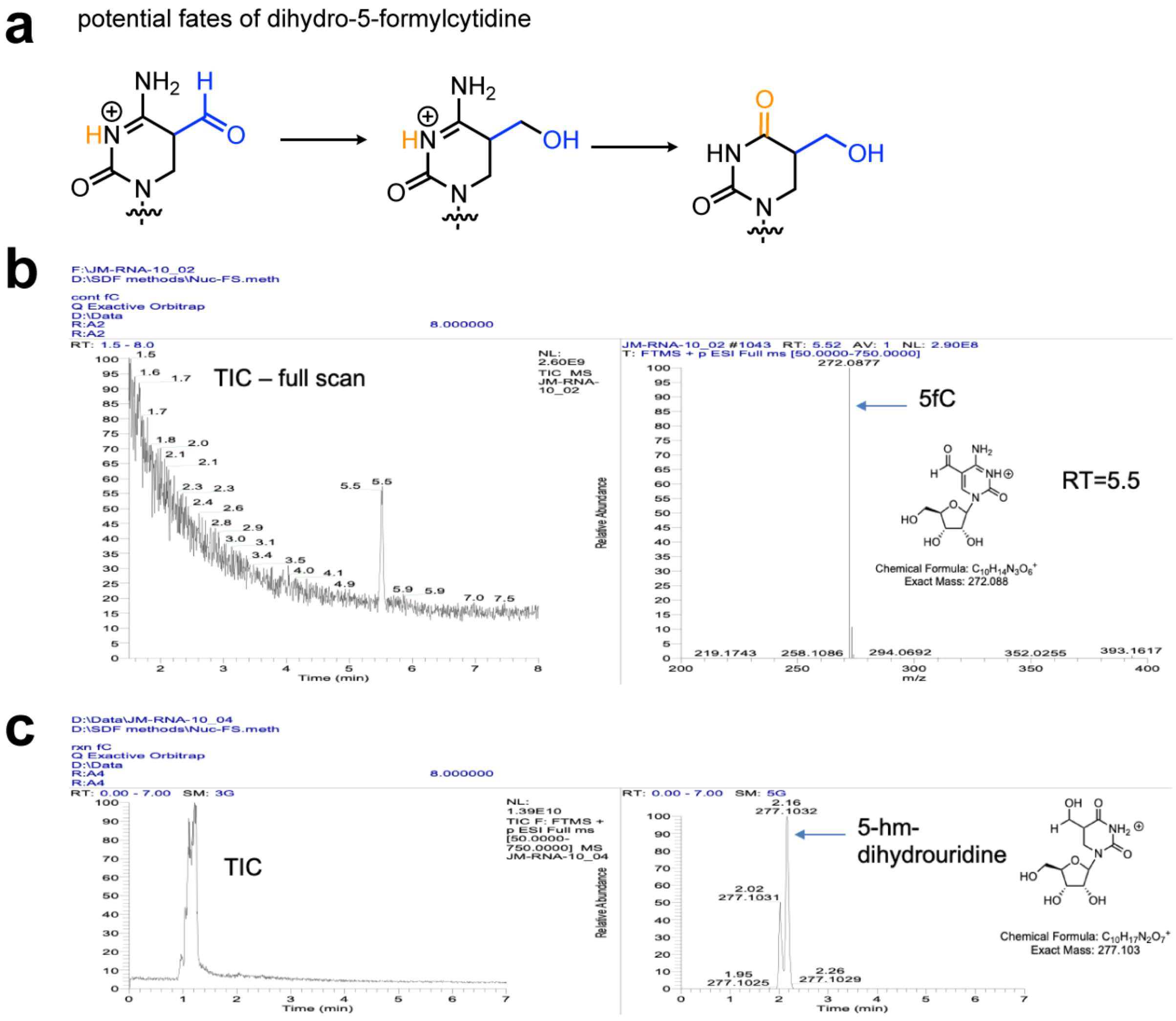


**Figure S4**. Product analysis of 5-formylcytidine following $NaCNBH_3$ (pH 1) treatment. (a) Potential reactions involved in conversion of 5-formylcytidine to 5-hydroxymethyl-dihydrocytidine due to $NaCNBH_3$ (pH 1) treatment. (b) LC-MS analysis of 5fC starting material. (c) LC-MS analysis indicating formation of 5-hydroxymethyl-dihydrocytidine (5-hm-dihydrouridine) from 5fC following $NaCNBH_3$ (pH 1) treatment.

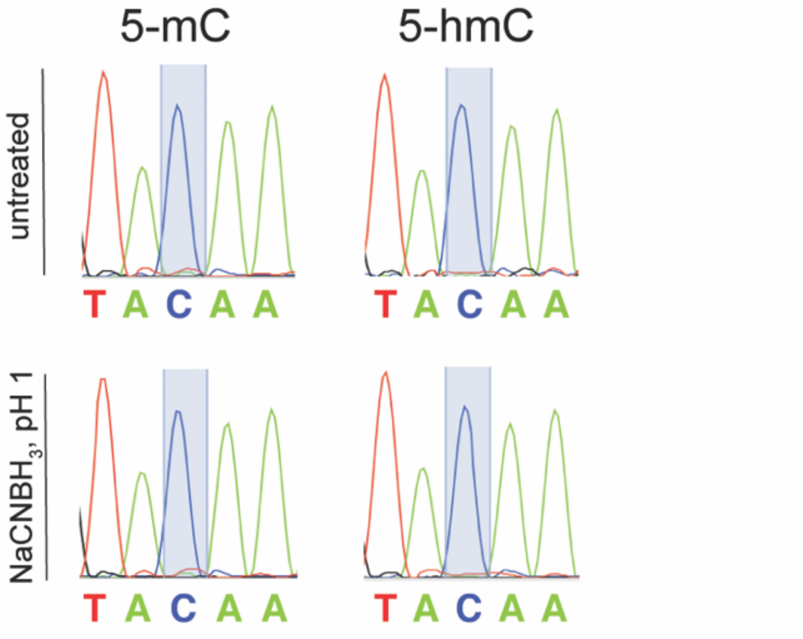


**Figure S5**. Sanger sequencing traces of 5mC and 5hmC sites in model RNAs following $NaCNBH_3$ treatment (pH 1) confirms the lack of reactivity of these modified nucleobases. Model RNAs correspond to 'single C' synthetic RNA used in Figure 3 of main text, transcribed with 5mCTP and 5hmCTP, respectively.

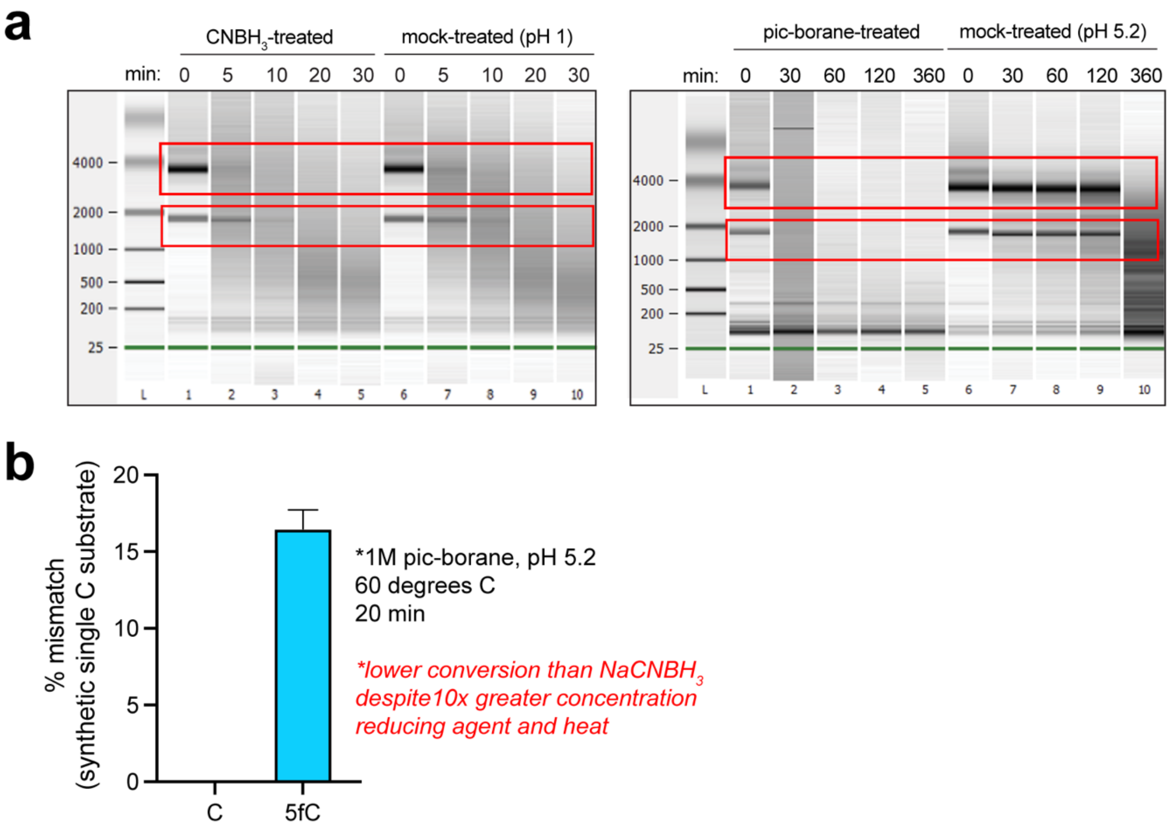


**Figure S6**. Assessing degradation as a funtion of pH. (a) Degradation of total RNA following treatment with $NaCNBH_3$ (100 mM, pH 1; left) or pic-borane (1M, pH 5.2; right). (b) Conversion of 5fC in model RNA after treatment with pic-borane for 20 min (1M, 60 degrees).

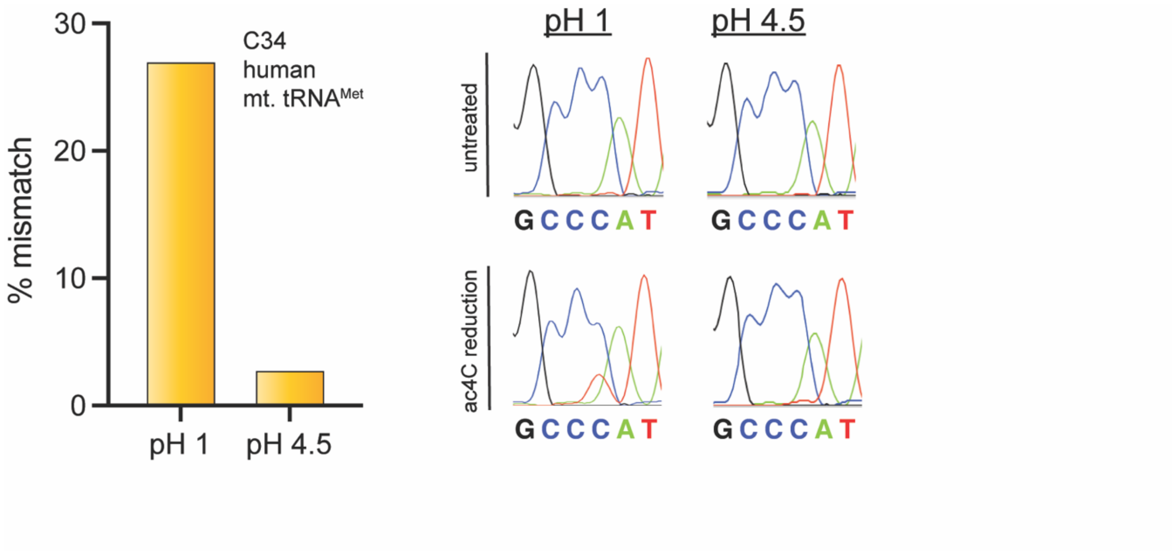


**Figure S7**. Misincorporation signal for 5fC in mitochondrial $tRNA^{Met}$ is dependent on pH of $NaCNBH_3$ sequencing reaction.

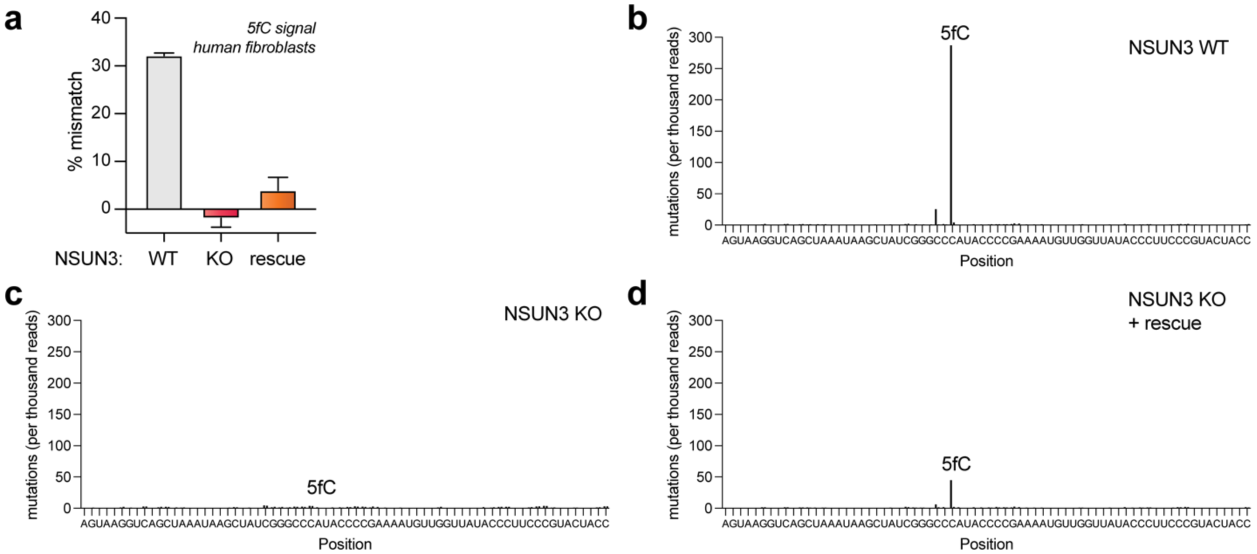


**Figure S8**. Applying a protonation-dependent sequencing reaction to detect disease-associated changes in 5fC levels within human tRNA$^{Met}$ (a) Sanger sequencing results indicating % misincorporation at C34 in human tRNA$^{Met}$ from wild-type (left), NSUN3 KO (middle), and NSUN3 KO/rescued (right) fibroblasts. Values indicate the C->T misincorporation percentage observed in $NaCNBH_3$ (pH 1) treated samples minus controls. (b) Next-generation sequencing analysis of $NaCNBH_3$ (pH 1)-dependent misincorporations in wild-type fibroblasts. (c) Next-generation sequencing analysis of $NaCNBH_3$ (pH 1)-dependent misincorporations in wild-type fibroblasts. (d) Next-generation sequencing analysis of $NaCNBH_3$ (pH 1)-dependent misincorporations in wild-type fibroblasts. All misincorporations at C34 were C->T transversions and were normalized to overall sequencing depth using ShapeMapper2.[19]

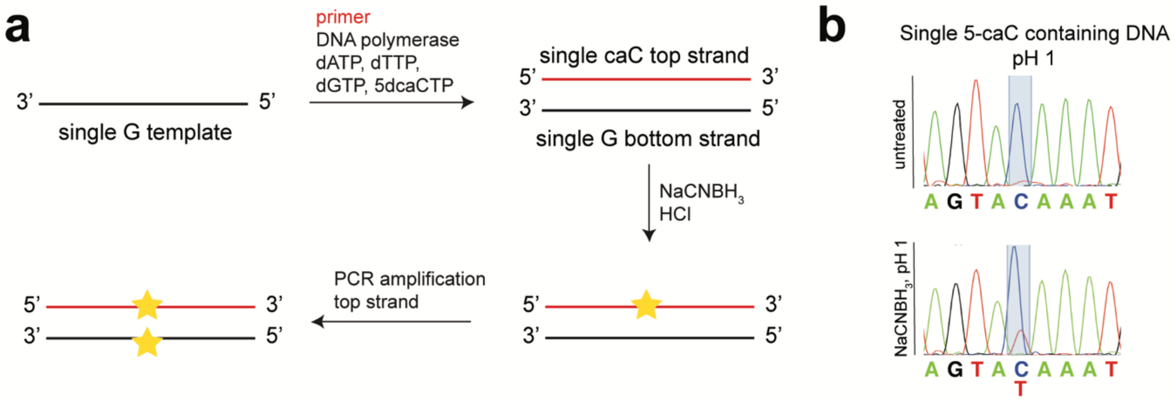


**Figure S9**. Assessing a protonation-dependent sequencing reaction for detection of oxidized cytidine residues in DNA. (a) A model DNA substrate containing a single 5caC site was produced using first-strand cDNA synthesis. This duplex was then subjected to $NaCNBH_3$ (pH 1) treatment, PCR-amplified, and analyzed by Sanger sequencing. (b) $NaCNBH_3$ (pH 1) treatment can generate partial misincorporations at 5caC sites in model DNA substrates.

### Experimental procedures and protocols

#### General materials and methods

Unless otherwise specified, all chemicals and solvents were purchased from Sigma, VWR, or Fisher and used without further purification. Modified nucleosides and nucleotides were purchased for Trilink Biotechnologies (San Diego, CA). Templates for in vitro transcription reactions and oligonucleotides for PCR amplification were purchased from Integrated DNA Technologies (Coralville, IA). TGIRT-III reverse transcriptase was purchased from Ingex. Analytical analyses of 5fC, 5caC, and their reduced products were performed using a Shimadzu LC-20AD coupled to a Thermo TSQ-ultra triple-quadrupole mass spectrometer as described previously.[1]

#### Computational chemistry

Quantum chemistry calculations were performed using a local version of the GAMESS[2,3] package where 6-31G(d) basis sets (spherical harmonics) were used[4,5]. Density functional theory (DFT)[6] utilizing the B3LYP[7–9] functional was used to optimize geometries and water solvent effects were included via the polarizable continuum model (PCM)[10–13] approach. Molecular orbitals were illustrated with MacMolPlt13.[14]

#### UV spectroscopic analysis of nucleoside reduction

Model reactions to assess the rate of reduction of C, 5mC, 5hmC, 5fC, d5fC, 5caC, 5ClC, and ac4C by NaCNBH3 were performed using the free nucleosides as described previously.[15] Briefly, stock solutions of $NaCNBH_3$ (1 M) and each modified nucleoside (2.5 mM) were prepared fresh daily in water. Reactions (100 µl) consisted of one of the aforementioned free nucleosides (100 µM), NaCNBH3 (100 mM), and buffer (100 mM). The buffer was selected based on the desired pH of the reaction; HCl was used for pH 1 reactions, sodium phosphate was used for pH 2.5 reactions, sodium acetate was used for pH 4.5 reactions, and sodium phosphate was used for pH 7.2 reactions. At the indicated time point, reactions were quenched with 30 µl of 1 M Tris-HCl (pH 8.0) and added to Greiner-UV Star 96-well microplates for analysis. Reduction of each modified nucleoside was analysed on a Biotek Synergy plate reader by monitoring the absorbance at *λ*max (*λ*max = 280 nm for 5fC, 5caC, and 5ClC; *λ*max = 300 nm for ac4C; and *λ*max = 270 nm for cytidine). The percentage decrease in each modified nucleoside was calculated from absorbance (*A*) values at *λ*max using the formula: Percentage decrease = $(A_{\text{nucleoside}}(\text{start}) - A_{\text{nucleoside}}(\text{end}))/(A_{\text{nucleoside}}(\text{untreated}) - A_{\text{water}}(\text{blank})) \times 100$.

#### LC-MS analysis of nucleosides

Model reactions to assess the reduction products of 5fC and 5caC by NaCNBH3 were performed using free 5-carboxycytidine and 5-formylcytidine nucleosides. A fresh stock solution of NaCNBH3 (1 M) was prepared daily in water. Reactions (100 µl) consisted of 5caC or 5fC (100 µM), NaCNBH3 (100 mM) and HCl (100 mM). After incubating for 30 minutes at room temperature, the reactions were quenched with 30 µl of 1 M Tris-HCl (pH 8.0) and stored on dry ice until LC-MS analysis. LC-MS analysis of free nucleosides was performed as described previously.[1,16]

**In vitro transcription of model RNAs containing a single C, 5mC, 5hmC, 5fC and 5caC**
In vitro transcription was performed with the HiScribe T7 Kit (NewEngland Biolabs), according to the manufacturer's instructions using DNA templates containing a T7 promoter upstream of a template sequence harboring a single cytidine as described previously.[15] To produce RNAs containing modified cytidines, CTP was replaced in the reaction mixture with 5mCTP, 5hmCTP, 5fCTP, or 5caCTP (5 mM), respectively. Synthesized RNA was mixed with 1X RNA denaturing RNA loading buffer and heated at 95 °C for 4 min, cooled on ice, loaded onto a 14% denaturing polyacrylamide gel, and run at 400 V (20 v/cm) for 5 h. Gel bands were excised, put in crush and soak buffer at 4 °C overnight, and desalted to yield pure RNA for reaction-based sequencing analyses.

**$NaCNBH_3$ reduction of RNA**
RNA samples were treated with sodium cyanoborohydride (100 mM in $H_2O$) or vehicle ($H_2O$) in a final reaction volume of 100 µL. Reactions were initiated by addition of 1 M HCl to a final concentration of 100 mM and incubated for 20 minutes at room temperature. Reactions were stopped by neutralizing the pH by the addition of 30 µL 1 M Tris-HCl pH 8.0. The quenched reactions were adjusted to 200 µL with H2O, purified via ethanol precipitation, and desalted via 70% ethanol wash. The pelleted RNA was dried using a Speedvac, resuspended in H2O, and quantified using a Nanodrop 2000 spectrophotometer. Samples were stored in -20 °C until reverse transcription was performed.

**Dot blot analysis of treated RNA**
To assess reduction of 5fC and 5caC in RNA, in vitro transcribed RNA harboring a single modification were treated with $NaCNBH_3$ or vehicle (water) and analyzed by dot blot. Briefly, purified RNA after the treatment was spotted onto Amersham Hybond-N+ membranes (GE Healthcare). Samples were crosslinked twice with 150 mJ/cm$^2$ in the UV254nm Stratalinker 2400 (Stratagene, San Diego, CA). Following crosslinking, samples were blocked with 5% non-fat milk in TBST at room temperature for 30 min, and then probed with anti-fC (1:5000) or anti-caC (1:1000) (Active Motif, Carlsbad, CA) antibody in 5% non-fat milk in TBST at 4 °C overnight. Membranes were washed three times with TBST, and incubated with HRP-conjugated secondary anti-rabbit IgG (1:2000 dilution, Cell Signaling) for 1 h. Membranes were washed four times with TBST and developed with the SuperSignal ELISA Femto Maximum Sensitivity Substrate (Pierce, ThermoScientific, Waltham, MA).

**2-picoline borane complex (pic-borane) reduction of RNA**
Reactions were carried out as previously described by Liu et al.[17] Briefly, RNA was treated with pic-borane (1.63 M in DMSO) or vehicle ($H_2O$) in a final reaction volume of 50 µL. Reactions were kept at pH 5.2 using sodium acetate buffer (300 mM) and were incubated at 70 °C and 850 r.p.m. in an Eppendorf ThermoMixer for 20 min. Reactions were immediately purified via ethanol precipitation and desalted via 70% ethanol wash. The pelleted RNA was dried using a Speedvac, resuspended in $H_2O$, and quantified using a Qubit 4 fluorimeter (Thermo Fisher Scientific). Samples were stored at -20 °C prior to reverse transcription and PCR.

**RT and PCR of model RNA substrates containing a single C or modified C**
Gel-purified RNA samples (~200 ng) were directly treated with a reducing agent ($NaCNBH_3$ or pic-borane) or water and purified as described above. For stoichiometry experiments only, the gel purified single 5fC and C-containing RNA were first mixed to produce 100, 90, 80, 60, 50, 40, 20, 10, and 0% 5fC:C ratios, then were subjected to $NaCNBH_3$ reduction and purification. Briefly, RNA samples from individual reactions (~200-500 pg) were incubated with rev (RT) primer for single C substrates (#3 from primer table below; 4.0 pmol) in a final volume of 20 µL. Individual reactions were heated to 65 °C for 3 min and transferred to ice for 1 min to facilitate annealing in

1X TGIRT reaction buffer (50 mM Tris-HCl (pH 8.3), 75 mM KCl and 3 mM $MgCl_2$). After annealing, reverse transcriptions were performed with the TGIRT-III (Ingex) enzyme by first adding 5 mM DTT, 25 units RNasin, 100 units TGIRT RT, and incubating 20 minutes at room temperature. Then 500 µM dNTPs (5 mM GTP, 10 mM CTP, ATP, and TTP) were added and incubated at 57 °C for 60 min. Reactions were quenched by increasing the temperature to 70 °C for 15 min. The cDNA products from the reactions and controls were directly used in PCR. PCR reactions were set up with 2 µL cDNA in 50 µL PCR reaction with Phusion Hot start flex (New England Biolabs). Reaction conditions: 1X supplied HF buffer, 2.5 pmole each forward and reverse primer (PCR primers for single C substrates, #4 and 5 from primer table below), 200 µM each dNTP, 2 units Phusion hot start enzyme, 2 µL template (Thermocycling conditions: 65.3 °C annealing, 34 cycles). PCR products were run on a 2% agarose gel, stained with SYBR safe, and visualized on UV transilluminator at 302 nm. Bands of the desired size were excised from the gel and DNA extracted using QIA-quick gel extraction kit from Zymo and submitted for Sanger sequencing (GeneWiz) using the “forward PCR primer for single C substrate”. Processed sequencing traces were viewed using 4Peaks software. Peak height for each base was measured and the percent misincorporation was determined using the equation: “Percent misincorporation = (peak intensity of T)/(sum of C and T base peaks)*100%”. Final misincorporation values were determined by subtracting background water control misincorporation levels from that of the corresponding reactions.

**Cell culture and total RNA isolation from human cell lines.**
HEK293 cells were grown in DMEM (Life Technologies) supplemented with 10% fetal bovine serum at 37 °C in a humidified atmosphere with 5% CO2. Patient-derived fibroblasts (NSUN3 KO and NSUN3 rescue) were identical to those reported previously[18] and were cultured in high glucose DMEM (Life Technologies), supplemented with 10% FBS, 1% penicillin/streptomycin, and 200 mM uridine at 37 °C and 5% $CO^2$. Cells were grown until ~80 - 90% confluency, detached by addition of Trypsin-EDTA and harvested by centrifugation (500 g, 2 min at room temperature). Cells were washed twice with PBS and pellets were stored frozen at -80 °C. Total RNA from human cells was extracted using TRIzol according to the manufacturer's protocol. 1 mL TRIzol was used per 1 x $10^7$ cells. The RNA pellet was resuspended in water and quantified by UV absorbance and stored at −80 °C. Typical extractions were carried out with 1 X $10^7$ cells and yielded 400 µg of total RNA. Isolated total RNA was incubated with Turbo DNase (Invitrogen) for 30 min at 37 °C to remove any DNA contamination prior to $NaCNBH_3$ treatment and purification as described above.

**5fC sequencing in human mitochondrial tRNA$^{met}$**

DNase-treated cellular total RNA samples (~5 µg) was subjected to $NaCNBH_3$ treatment and purification as described above. Treated RNA samples from individual reactions (~1-2 µg) were incubated with human mitochondrial mRNA$^{Met}$ RT rev primer (primer # 6 from table below; 4.0 pmol) in a final volume of 20 µL. Individual reactions were subjected to reverse transcription with TGIRT III enzyme as described above. The cDNA products from the reactions and controls were purified using G-25 columns and used for the first PCR reaction as follows:

Reaction conditions for PCR 1: Mix 1X Q5 reaction buffer, 0.5 µM each step 1 forward and step 1 reverse primer, 200 µM each dNTP, 1 unit Q5 hot start enzyme (New England Biolabs), 5 µL cDNA template, and water up to 50 µL in a PCR tube and use the following parameters for the thermocycler.

| Step | Denature | Anneal | Extend |
|---|---|---|---|
| 1 | 98 °C, 30 s | | |
| 2–6 | 98 °C, 10 s | Tm °C, 30 s | 72 °C, 20 s |
| 7 | | | 72 °C, 2 min |

PCR products from the above reaction were purified using Purelink PCR micro spin columns according to the manufacturer's instructions. The samples were eluted with 10 µL of water.

PCR 2 reactions were set up with 10 µL of the PCR 1 purified products as below.

Reaction conditions for PCR 2: Mix 1X Q5 reaction buffer, 0.5 µM each universal forward (with barcodes for Next Generation Sequencing) and universal reverse primer, 200 µM each dNTP, 1 unit Q5 hot start enzyme, 10 µL cDNA template, and water up to 50 µL in a PCR tube and use the following parameters for the thermocycler.

| Step | Denature | Anneal | Extend |
|---|---|---|---|
| 1 | 98 °C, 30 s | | |
| 2–26 | 98 °C, 10 s | 65 °C, 30 s | 72 °C, 20 s |
| 27 | | | 72 °C, 2 min |

PCR products were run on a 2% agarose gel, stained with SYBR safe and visualized on UV transilluminator at 302 nm. Bands of the desired size were excised from the gel and DNA extracted using Zymoclean gel DNA recovery kit and pooled to make a library of 4 nM. Libraries were analyzed using a MiSeq Benchtop Sequencing system using the MiSeq Nano 300 kit (2x150 bp format). Misincorporations across the tRNA$^{Met}$ sequence were counted and normalized to overall sequencing depth using the ShapeMapper2 software.[19]

**DNA templates and primers**

| | | |
|---|---|---|
| 1 | *"Single ac4C" in vitro transcription template (red = single C)* | 5'-GGGAGAGAAGGTTGGAAGAGATGTTAGTACAAATGATTGTGTAGAGGGAGAAAAGTGATGAGTGAAGTTGTGATGAGAAGAGGGAGAGATGTTAAAGATGGGAAGGTGATAAGTGTTTGTAGAGGAAGAGAATGTATTAATGGTGGGAG-3' |
| 2 | *"Single 5fC/5caC" RNA product (red = single fC/caC)* | 5'-GGGAGAGAAGGUUGGAAGAGAUGUUAGUACAAAUGAUUGUGUAGAGGGAGAAAAGUGAUGAGUGAAGUUGUGAUGAGAAGAGGGAGAGAUGUUAAAGAUGGGAAGGUGAUAAGUGUUUGUAGAGGAAGAGAAUGUAUUAAUGGUGGGAG-3' |
| 3 | IVT rev (RT primer) for single C substrate | 5'-CTCCCACCATTAATACATTCTCTTCCTC-3' |
| 4 | IVT rev (PCR primer) for single C substrate | 5'-CCACCATTAATACATTCTCTTCCTCTACA-3' |
| 5 | IVT forward (PCR primer) for single C substrate | 5'-GGGAGAGAAGGTTGGAAGAGATGTTA-3' |
| 6 | human mt. mRNA-met RT rev primer | 5'-TGGTAGTACGGGAAGGG-3' |
| 7 | PCR 1 forward primer | 5'-GACTGGAGTTCAGACGTGTGCTCTTCCGATCTGTACGAGTAAGGTCAGCTAAATAAGC-3' |
| 8 | PCR 1 rev primer | 5'-CCCTACACGACGCTCTTCCGATCTGTACGTGGTAGTACGGGAAGGG-3' |
| 9 | universal forward primer | 5'-CAAGCAGAAGACGGCATACGAGAT[barcode]GTGACTGGAGTTCAGAC-3' |
| 10 | universal reverse primer | 5'-AATGATACGGCGACCACCGAGATCTACACTCTTTCCCTACACGACGCT |

| | | |
|---|---|---|
| | | CTTCCG-3’ |
| 11 | human mt. tRNA-met sanger sequencing primer | 5’-GACTGGAGTTCAGACGTG -3’ |

**Raw computational data:**

**I. LUMO energies (eV) and optimized structures (Å) of neutral systems**

**a) C**

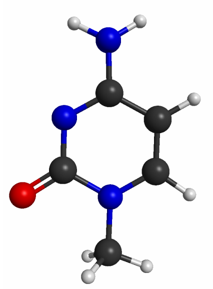

*E(LUMO) = -0.659*

| ATOM | CHARGE | X | Y | Z |
|---|---|---|---|---|
| C | 6.0 | -0.1674759148 | 0.4505095588 | 0.1010624572 |
| C | 6.0 | 1.0928964443 | -0.2206202887 | -0.0096166629 |
| C | 6.0 | 1.0542849288 | -1.5807159388 | -0.0583350746 |
| N | 7.0 | -0.1239509550 | -2.2575320426 | -0.0015235082 |
| C | 6.0 | -1.3580091696 | -1.5619572489 | 0.1112322059 |
| N | 7.0 | -1.3288043415 | -0.1981641816 | 0.1555160646 |
| H | 1.0 | 1.9503940339 | -2.1860373657 | -0.1451238501 |
| C | 6.0 | -0.1727681043 | -3.7194314105 | -0.0575208298 |
| H | 1.0 | 0.8431590692 | -4.1027280403 | -0.1659165142 |
| H | 1.0 | -0.7799496217 | -4.0406144392 | -0.9074272092 |
| H | 1.0 | -0.6191423772 | -4.1165991738 | 0.8579694235 |
| O | 8.0 | -2.4008750271 | -2.2252290600 | 0.1599100588 |
| N | 7.0 | -0.2030246223 | 1.8041250418 | 0.1879552683 |
| H | 1.0 | 0.6112189569 | 2.3512588655 | -0.0540014064 |
| H | 1.0 | -1.1029730208 | 2.2588540689 | 0.1062591561 |
| H | 1.0 | 2.0317484510 | 0.3173269451 | -0.0516034289 |

**b) m4C**

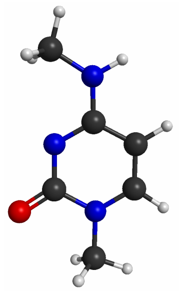

*E(LUMO) = -0.542*

| ATOM | CHARGE | X | Y | Z |
|---|---|---|---|---|
| C | 6.0 | -0.1833840376 | 0.4236638927 | 0.2600072220 |
| C | 6.0 | 1.0829133695 | -0.2348245062 | 0.0996844370 |
| C | 6.0 | 1.0516650250 | -1.5859962210 | -0.0452820668 |
| N | 7.0 | -0.1254123308 | -2.2725151626 | -0.0367412328 |
| C | 6.0 | -1.3613112064 | -1.5928117312 | 0.1251264480 |
| N | 7.0 | -1.3418521761 | -0.2357465796 | 0.2699386609 |
| H | 1.0 | 1.9503481344 | -2.1797605454 | -0.1746566547 |
| C | 6.0 | -0.1646781281 | -3.7268115725 | -0.1937562410 |
| H | 1.0 | 0.8548424086 | -4.0951759794 | -0.3199132192 |
| H | 1.0 | -0.7620914872 | -3.9944344978 | -1.0690748229 |
| H | 1.0 | -0.6152605187 | -4.1896123844 | 0.6882441406 |
| O | 8.0 | -2.4013039906 | -2.2638542459 | 0.1262240519 |
| N | 7.0 | -0.2141450617 | 1.7665517626 | 0.4089116133 |
| H | 1.0 | 0.6640582647 | 2.2643828766 | 0.3874868043 |
| H | 1.0 | 2.0191578851 | 0.3099525336 | 0.0923098393 |
| C | 6.0 | -1.4386193504 | 2.5333305138 | 0.5668730731 |
| H | 1.0 | -2.0000227078 | 2.2032973053 | 1.4477978879 |
| H | 1.0 | -2.0900614587 | 2.4299677289 | -0.3086328426 |
| H | 1.0 | -1.1704506932 | 3.5843961426 | 0.6885315518 |

**c) 5mC**

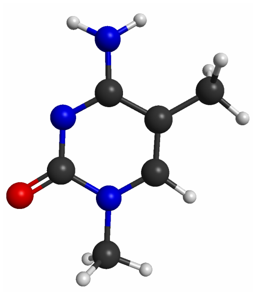

*E(LUMO) = -0.618*

| ATOM | CHARGE | X | Y | Z |
|---|---|---|---|---|
| C | 6.0 | -0.1640922638 | 0.4434115240 | 0.1024389120 |
| C | 6.0 | 1.1113432268 | -0.2182304742 | -0.0066136534 |
| C | 6.0 | 1.0477428348 | -1.5797131338 | -0.0553767775 |
| N | 7.0 | -0.1329555763 | -2.2573645589 | -0.0009933484 |
| C | 6.0 | -1.3626669529 | -1.5656806787 | 0.1119906996 |
| N | 7.0 | -1.3256469304 | -0.2023992271 | 0.1556482040 |
| H | 1.0 | 1.9409344752 | -2.1907569578 | -0.1403304442 |
| C | 6.0 | 2.4132159637 | 0.5362699478 | -0.0585588776 |
| H | 1.0 | 3.2577752729 | -0.1551584832 | -0.1362348418 |
| H | 1.0 | 2.4609329321 | 1.2113359694 | -0.9241900534 |
| H | 1.0 | 2.5654676541 | 1.1470925059 | 0.8412814454 |
| C | 6.0 | -0.1782845137 | -3.7192157515 | -0.0591169179 |
| H | 1.0 | 0.8381335214 | -4.1005063673 | -0.1697179558 |
| H | 1.0 | -0.7859518024 | -4.0407146692 | -0.9086796855 |
| H | 1.0 | -0.6224323250 | -4.1193364033 | 0.8563069713 |
| O | 8.0 | -2.4099131994 | -2.2244440293 | 0.1607917281 |
| N | 7.0 | -0.2068795599 | 1.7986575949 | 0.1917268477 |
| H | 1.0 | 0.5933837305 | 2.3556513945 | -0.0712352364 |
| H | 1.0 | -1.1133879774 | 2.2407900180 | 0.1108629237 |

**d) 5hmC**

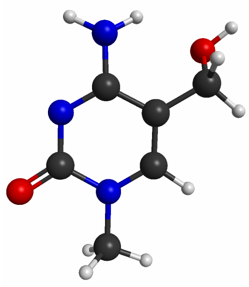

*E(LUMO) = -0.686*

| ATOM | CHARGE | X | Y | Z |
|---|---|---|---|---|
| C | 6.0 | -0.2186021993 | 0.4557703917 | 0.2237607898 |
| C | 6.0 | 1.0664400699 | -0.1798427394 | 0.0656977937 |
| C | 6.0 | 1.0347437428 | -1.5355426618 | -0.0785491505 |
| N | 7.0 | -0.1287249502 | -2.2371679160 | -0.0424897840 |
| C | 6.0 | -1.3698515243 | -1.5786813271 | 0.1604168467 |
| N | 7.0 | -1.3630022877 | -0.2178821002 | 0.2712138899 |
| H | 1.0 | 1.9396549821 | -2.1191193515 | -0.2173115254 |
| C | 6.0 | 2.3555067160 | 0.5875456190 | 0.1302267894 |
| C | 6.0 | -0.1488684269 | -3.6936943012 | -0.1922454349 |
| H | 1.0 | 0.8726213064 | -4.0465647094 | -0.3426754151 |
| H | 1.0 | -0.7639704661 | -3.9730155699 | -1.0513300111 |
| H | 1.0 | -0.5707600281 | -4.1567350361 | 0.7032690726 |
| O | 8.0 | -2.3972554845 | -2.2646976325 | 0.2125860977 |
| N | 7.0 | -0.2638720434 | 1.8070490322 | 0.3611657659 |
| H | 1.0 | 0.5182338462 | 2.3289769331 | -0.0220769159 |
| H | 1.0 | -1.1786054971 | 2.2380069126 | 0.3057914895 |
| H | 1.0 | 3.1990008969 | -0.0928001414 | -0.0491430209 |
| H | 1.0 | 2.4854736905 | 1.0222224996 | 1.1327600566 |
| O | 8.0 | 2.3327194164 | 1.6403083917 | -0.8505785637 |
| H | 1.0 | 3.0821505705 | 2.2293994665 | -0.6614526904 |

**e) ac4C**

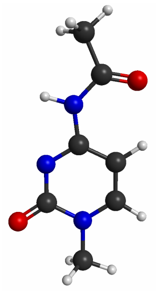

*E(LUMO) = -1.363*

| ATOM | CHARGE | X | Y | Z |
|---|---|---|---|---|
| C | 6.0 | 0.0741238896 | 0.4130928322 | 0.1728833994 |
| C | 6.0 | 1.2692864659 | -0.3498564920 | 0.0790523524 |
| C | 6.0 | 1.1030826866 | -1.7045234992 | -0.0082318484 |
| N | 7.0 | -0.1264706481 | -2.2732435249 | -0.0061259544 |
| C | 6.0 | -1.2983931103 | -1.4770605608 | 0.0835055750 |
| N | 7.0 | -1.1382523962 | -0.1203893552 | 0.1749607277 |
| H | 1.0 | 1.9450027812 | -2.3842108631 | -0.0830405738 |
| C | 6.0 | -0.3033423538 | -3.7254154169 | -0.0977439147 |
| H | 1.0 | 0.6792875004 | -4.1973531484 | -0.1306147860 |
| H | 1.0 | -0.8661362145 | -3.9748544727 | -1.0006174509 |
| H | 1.0 | -0.8596972920 | -4.0836929684 | 0.7712391702 |
| O | 8.0 | -2.3981141060 | -2.0355985448 | 0.0761198457 |
| N | 7.0 | 0.0741608382 | 1.8014198699 | 0.2717123381 |
| H | 1.0 | -0.8608393819 | 2.1908502394 | 0.3349563065 |
| H | 1.0 | 2.2435460355 | 0.1106846875 | 0.0783109216 |
| C | 6.0 | 1.1414483210 | 2.6876639143 | 0.3053938526 |
| O | 8.0 | 2.3122357239 | 2.3356371172 | 0.2326911966 |
| C | 6.0 | 0.7222395217 | 4.1360317819 | 0.4614867983 |
| H | 1.0 | 1.5889503729 | 4.7748391050 | 0.2858153637 |
| H | 1.0 | -0.0784055340 | 4.4019324772 | -0.2369166204 |
| H | 1.0 | 0.3522744997 | 4.3103112318 | 1.4791573608 |

**f) 5fC**

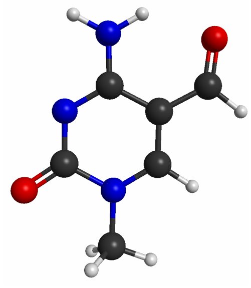

*E(LUMO) = -1.548*

| ATOM | CHARGE | X | Y | Z |
|---|---|---|---|---|
| C | 6.0 | -0.2027888206 | 0.4775898862 | 0.0743706372 |
| C | 6.0 | 1.0824704356 | -0.1915058410 | -0.0202271389 |
| C | 6.0 | 1.0435937330 | -1.5704126940 | -0.0559290291 |
| N | 7.0 | -0.1088849129 | -2.2475935137 | -0.0052813855 |
| C | 6.0 | -1.3684176087 | -1.5497029706 | 0.0918384850 |
| N | 7.0 | -1.3519086448 | -0.1917936596 | 0.1265920829 |
| H | 1.0 | 1.9547904588 | -2.1577669097 | -0.1278596942 |
| C | 6.0 | 2.3623180209 | 0.4846332493 | -0.0810219374 |
| H | 1.0 | 3.2385634548 | -0.1890997117 | -0.1511474857 |
| C | 6.0 | -0.1507780630 | -3.7133260292 | -0.0461399043 |
| H | 1.0 | 0.8677987225 | -4.0917580192 | -0.1413366165 |
| H | 1.0 | -0.7502732694 | -4.0392514396 | -0.8985212511 |
| H | 1.0 | -0.6047265437 | -4.0955623283 | 0.8709010253 |
| O | 8.0 | -2.3900539263 | -2.2337422250 | 0.1370844678 |
| N | 7.0 | -0.2371106409 | 1.8166424199 | 0.1108198094 |
| H | 1.0 | 0.6355001135 | 2.3333942585 | 0.0700992269 |
| H | 1.0 | -1.1283219075 | 2.2901208286 | 0.1729422317 |
| O | 8.0 | 2.5283202187 | 1.7066325992 | -0.0609730436 |

**g) 5caC**

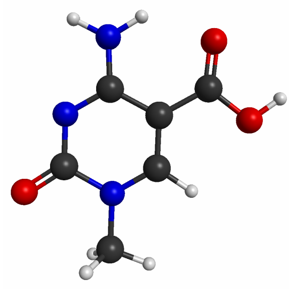

*E(LUMO) = -1.309*

| ATOM | CHARGE | X | Y | Z |
|---|---|---|---|---|
| C | 6.0 | -0.1556294385 | 0.4740214977 | 0.0740314018 |
| C | 6.0 | 1.1218127432 | -0.2092024653 | -0.0057277209 |
| C | 6.0 | 1.0664574811 | -1.5859153949 | -0.0411698226 |
| N | 7.0 | -0.0961621606 | -2.2514382582 | -0.0033608139 |
| C | 6.0 | -1.3457604385 | -1.5443969867 | 0.0783325107 |
| N | 7.0 | -1.3113480064 | -0.1880992655 | 0.1126767469 |
| H | 1.0 | 1.9686351406 | -2.1833750634 | -0.1021715800 |
| C | 6.0 | 2.4023002918 | 0.4979409092 | -0.0507499950 |
| C | 6.0 | -0.1496098313 | -3.7164981748 | -0.0446710851 |
| H | 1.0 | 0.8667230179 | -4.1042322681 | -0.1254000893 |
| H | 1.0 | -0.7400584201 | -4.0381160545 | -0.9051223437 |
| H | 1.0 | -0.6199630309 | -4.0949712818 | 0.8657330202 |
| O | 8.0 | -2.3781624709 | -2.2152725708 | 0.1116709944 |
| N | 7.0 | -0.2031900072 | 1.8138632397 | 0.1119840476 |
| H | 1.0 | 0.6583676881 | 2.3466191574 | 0.0804417387 |
| H | 1.0 | -1.1034751308 | 2.2706106992 | 0.1624346724 |
| O | 8.0 | 2.5358729971 | 1.7189616997 | -0.0278400421 |
| O | 8.0 | 3.4697136754 | -0.3268394412 | -0.1225898567 |

| | | | | |
|---|---|---|---|---|
| H | 1.0 | 4.2674280901 | 0.2363022523 | -0.1494529134 |

**h) 5-ClC**

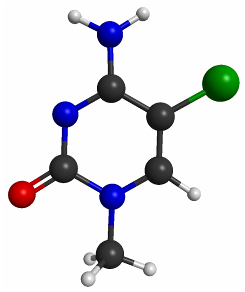

*E(LUMO) = -0.955*

| ATOM | CHARGE | X | Y | Z |
|---|---|---|---|---|
| C | 6.0 | -0.1704085200 | 0.4672051160 | 0.0877682588 |
| C | 6.0 | 1.0835586614 | -0.2276736078 | -0.0187871683 |
| C | 6.0 | 1.0540529596 | -1.5891872599 | -0.0619504827 |
| N | 7.0 | -0.1224008308 | -2.2606449599 | -0.0036475265 |
| C | 6.0 | -1.3568916528 | -1.5616770444 | 0.1041952664 |
| N | 7.0 | -1.3198986210 | -0.1986211876 | 0.1423696409 |
| H | 1.0 | 1.9575287824 | -2.1825350312 | -0.1443059295 |
| C | 6.0 | -0.1697045695 | -3.7247054759 | -0.0521516052 |
| H | 1.0 | 0.8462138972 | -4.1081936973 | -0.1559652877 |
| H | 1.0 | -0.7753000786 | -4.0467332010 | -0.9024194840 |
| H | 1.0 | -0.6186678238 | -4.1132626602 | 0.8653201069 |
| O | 8.0 | -2.3995086981 | -2.2206529640 | 0.1548321748 |
| N | 7.0 | -0.2099587361 | 1.8127355409 | 0.1570246440 |
| H | 1.0 | 0.6120661968 | 2.3744980011 | -0.0112789229 |
| H | 1.0 | -1.1130706801 | 2.2667209478 | 0.1334926154 |

| | | | | |
|---|---|---|---|---|
| CL | 17.0 | 2.6089954834 | 0.6339323835 | -0.0930514803 |

### II. LUMO energies (eV) and optimized structures (Å) of protonated systems

### a) C-H+

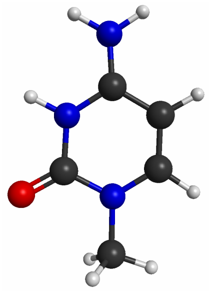

*E(LUMO) = -2.008*

| ATOM | CHARGE | X | Y | Z |
|---|---|---|---|---|
| C | 6.0 | -0.1674759148 | 0.4505095588 | 0.1010624572 |
| C | 6.0 | 1.0928964443 | -0.2206202887 | -0.0096166629 |
| C | 6.0 | 1.0542849288 | -1.5807159388 | -0.0583350746 |
| N | 7.0 | -0.1239509550 | -2.2575320426 | -0.0015235082 |
| C | 6.0 | -1.3580091696 | -1.5619572489 | 0.1112322059 |
| N | 7.0 | -1.3288043415 | -0.1981641816 | 0.1555160646 |
| H | 1.0 | 1.9503940339 | -2.1860373657 | -0.1451238501 |
| C | 6.0 | -0.1727681043 | -3.7194314105 | -0.0575208298 |
| H | 1.0 | 0.8431590692 | -4.1027280403 | -0.1659165142 |
| H | 1.0 | -0.7799496217 | -4.0406144392 | -0.9074272092 |
| H | 1.0 | -0.6191423772 | -4.1165991738 | 0.8579694235 |

| | | | | |
|---|---|---|---|---|
| O | 8.0 | -2.4008750271 | -2.2252290600 | 0.1599100588 |
| N | 7.0 | -0.2030246223 | 1.8041250418 | 0.1879552683 |
| H | 1.0 | 0.6112189569 | 2.3512588655 | -0.0540014064 |
| H | 1.0 | -1.1029730208 | 2.2588540689 | 0.1062591561 |
| H | 1.0 | 2.0317484510 | 0.3173269451 | -0.0516034289 |

**b) m4C-H+**

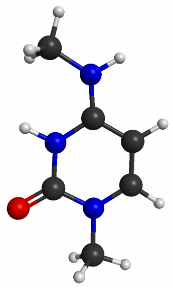

*E(LUMO) = -1.932*

| ATOM | CHARGE | X | Y | Z |
|---|---|---|---|---|
| C | 6.0 | -0.1549869238 | 0.4823844406 | 0.2142450619 |
| C | 6.0 | 1.0663294740 | -0.2174682514 | 0.0157046843 |
| C | 6.0 | 1.0219082422 | -1.5747025136 | -0.0893972261 |
| N | 7.0 | -0.1385021964 | -2.2875549143 | -0.0123174423 |
| C | 6.0 | -1.3639368785 | -1.6593628148 | 0.1840660588 |
| N | 7.0 | -1.2919423544 | -0.2652427965 | 0.2892969193 |
| H | 1.0 | 1.9199658442 | -2.1609576484 | -0.2433498605 |
| C | 6.0 | -0.1601551765 | -3.7549533382 | -0.1315337257 |
| H | 1.0 | 0.8607178012 | -4.1005079294 | -0.2905478067 |
| H | 1.0 | -0.7861548643 | -4.0442122918 | -0.9779819270 |
| H | 1.0 | -0.5631662755 | -4.1915666674 | 0.7843950794 |

| | | | | |
|---|---|---|---|---|
| O | 8.0 | -2.4227088530 | -2.2544130619 | 0.2607288866 |
| N | 7.0 | -0.2252357420 | 1.8011249100 | 0.3243866435 |
| H | 1.0 | 0.6504885331 | 2.3072589312 | 0.2710223384 |
| H | 1.0 | 2.0029702276 | 0.3185289787 | -0.0525177212 |
| C | 6.0 | -1.4505847630 | 2.5723665616 | 0.5286264676 |
| H | 1.0 | -2.1538184568 | 2.4114934879 | -0.2954487879 |
| H | 1.0 | -1.1808356183 | 3.6272211329 | 0.5533773457 |
| H | 1.0 | -1.9246713005 | 2.3089486724 | 1.4802491711 |
| H | 1.0 | -2.1944233894 | 0.1776179922 | 0.4344027907 |

**c) 5mC-H+**

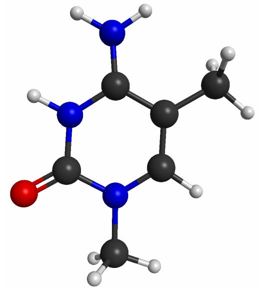

*E(LUMO) = -1.967*

| ATOM | CHARGE | X | Y | Z |
|---|---|---|---|---|
| C | 6.0 | -0.1439536598 | 0.4978685107 | 0.0529538465 |
| C | 6.0 | 1.0967283269 | -0.2029617069 | -0.0212341212 |
| C | 6.0 | 1.0174559124 | -1.5679880306 | -0.0506936206 |
| N | 7.0 | -0.1522357992 | -2.2678137568 | -0.0119503640 |
| C | 6.0 | -1.3811926565 | -1.6252068260 | 0.0666158658 |
| N | 7.0 | -1.2914135258 | -0.2312763216 | 0.0919823185 |
| H | 1.0 | 1.9143188424 | -2.1741808720 | -0.1084561332 |
| C | 6.0 | 2.4057710242 | 0.5381144378 | -0.0696445786 |
| H | 1.0 | 3.2385359680 | -0.1675935204 | -0.1197409554 |
| H | 1.0 | 2.4669368406 | 1.1883076236 | -0.9511422960 |

| | | | | |
|---|---|---|---|---|
| H | 1.0 | 2.5506270074 | 1.1594509008 | 0.8229142331 |
| C | 6.0 | -0.1776575396 | -3.7396206243 | -0.0490636299 |
| H | 1.0 | 0.8448028395 | -4.0984275618 | -0.1607257717 |
| H | 1.0 | -0.7828268442 | -4.0710823098 | -0.8948479099 |
| H | 1.0 | -0.6079033022 | -4.1239488167 | 0.8782775545 |
| O | 8.0 | -2.4505022672 | -2.2063378175 | 0.1085505052 |
| N | 7.0 | -0.2249036777 | 1.8214678299 | 0.0858265229 |
| H | 1.0 | 0.6111035097 | 2.3902664457 | 0.0554847455 |
| H | 1.0 | -1.1110513106 | 2.3102853072 | 0.1315775799 |
| H | 1.0 | -2.1911901882 | 0.2392903587 | 0.1433161483 |

**d) 5hmC-H+**

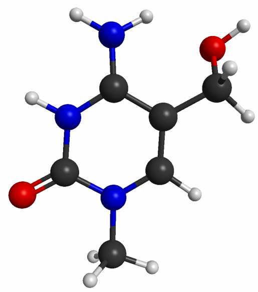

*E(LUMO) = -1.989*

| ATOM | CHARGE | X | Y | Z |
|---|---|---|---|---|
| C | 6.0 | -0.1933399409 | 0.5165715370 | 0.1596980357 |
| C | 6.0 | 1.0561828815 | -0.1653587442 | 0.0294934515 |
| C | 6.0 | 1.0053188834 | -1.5261707151 | -0.0804956121 |
| N | 7.0 | -0.1490004791 | -2.2476608270 | -0.0400816290 |
| C | 6.0 | -1.3855392080 | -1.6325306976 | 0.1280516311 |
| N | 7.0 | -1.3230149885 | -0.2388895926 | 0.2128826406 |
| H | 1.0 | 1.9116294126 | -2.1098752406 | -0.1973656468 |
| C | 6.0 | 2.3636132121 | 0.5807646760 | 0.1112356999 |
| C | 6.0 | -0.1532227350 | -3.7154471057 | -0.1632514721 |

| | | | | |
|---|---|---|---|---|
| H | 1.0 | 0.8724826725 | -4.0499064215 | -0.3140619129 |
| H | 1.0 | -0.7686656817 | -4.0074729660 | -1.0162390599 |
| H | 1.0 | -0.5606809947 | -4.1581612651 | 0.7478131400 |
| O | 8.0 | -2.4394335800 | -2.2380419653 | 0.1856231660 |
| N | 7.0 | -0.2774826198 | 1.8358386600 | 0.2369460865 |
| H | 1.0 | 0.5595958279 | 2.3655928589 | -0.0060496532 |
| H | 1.0 | -1.1670445121 | 2.3181027679 | 0.2881535159 |
| H | 1.0 | 3.1817153132 | -0.0911111928 | -0.1763870787 |
| H | 1.0 | 2.5397771897 | 0.8992527121 | 1.1486112013 |
| O | 8.0 | 2.3018545363 | 1.7230554153 | -0.7515383987 |
| H | 1.0 | 3.0646285583 | 2.2897068891 | -0.5472562391 |
| H | 1.0 | -2.2304684277 | 0.2115182473 | 0.3000700940 |

**e) m3C-H+**

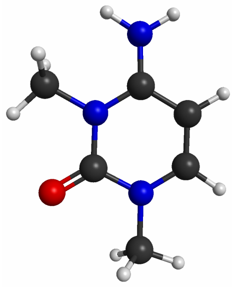

*E(LUMO) = -1.929*

| ATOM | CHARGE | X | Y | Z |
|---|---|---|---|---|
| C | 6.0 | -0.1490015073 | 0.5068135420 | 0.0590538208 |
| C | 6.0 | 1.0723959052 | -0.2070781291 | -0.0490431831 |
| C | 6.0 | 1.0261379294 | -1.5664343478 | -0.0740124039 |
| N | 7.0 | -0.1451899755 | -2.2524951324 | 0.0042660811 |
| C | 6.0 | -1.3715993424 | -1.6054631342 | 0.1179743202 |
| N | 7.0 | -1.3182687697 | -0.1941988840 | 0.1395840095 |
| H | 1.0 | 1.9223167729 | -2.1692715030 | -0.1586664820 |

| | | | | |
|---|---|---|---|---|
| C | 6.0 | -0.1831381357 | -3.7248533845 | -0.0249150670 |
| H | 1.0 | 0.8355641843 | -4.0898518781 | -0.1508030314 |
| H | 1.0 | -0.8038354992 | -4.0570559626 | -0.8589725138 |
| H | 1.0 | -0.6012540813 | -4.1010901345 | 0.9109855242 |
| O | 8.0 | -2.4220111807 | -2.2184260237 | 0.1926963983 |
| N | 7.0 | -0.1758650845 | 1.8353037254 | 0.0832362023 |
| H | 1.0 | 0.6909585939 | 2.3551070680 | 0.0270105144 |
| H | 1.0 | -1.0278691887 | 2.3746261358 | 0.1549727025 |
| H | 1.0 | 2.0117605898 | 0.3241981891 | -0.1156798754 |
| C | 6.0 | -2.6006837730 | 0.5224704742 | 0.2564150413 |
| H | 1.0 | -2.7505194333 | 1.1572382867 | -0.6213749772 |
| H | 1.0 | -2.6066228354 | 1.1215989942 | 1.1710887667 |
| H | 1.0 | -3.3904727585 | -0.2218453315 | 0.3041714124 |

**f) ac4C-H+**

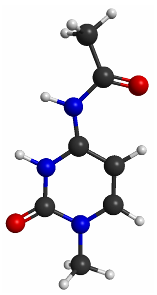

*E(LUMO) = -2.675*

| ATOM | CHARGE | X | Y | Z |
|---|---|---|---|---|
| C | 6.0 | 0.0697212300 | 0.4628759145 | 0.1897623796 |
| C | 6.0 | 1.2263494192 | -0.3304352405 | 0.0972576774 |
| C | 6.0 | 1.0592857375 | -1.6913853341 | 0.0034333312 |
| N | 7.0 | -0.1504184974 | -2.2931710365 | -0.0054037884 |
| C | 6.0 | -1.3333293186 | -1.5517268468 | 0.0795231239 |

| | | | | |
|---|---|---|---|---|
| N | 7.0 | -1.1327143653 | -0.1720726785 | 0.1787761069 |
| H | 1.0 | 1.9141974415 | -2.3533307800 | -0.0714031161 |
| C | 6.0 | -0.2992026245 | -3.7562100194 | -0.1078406774 |
| H | 1.0 | 0.6950395763 | -4.1987586336 | -0.1468535095 |
| H | 1.0 | -0.8567054937 | -4.0000530981 | -1.0140844126 |
| H | 1.0 | -0.8415627187 | -4.1240864468 | 0.7648215540 |
| O | 8.0 | -2.4404185149 | -2.0499647086 | 0.0706685352 |
| N | 7.0 | 0.0349500588 | 1.8167959985 | 0.2896418886 |
| H | 1.0 | -0.8823999078 | 2.2481831192 | 0.3518523063 |
| H | 1.0 | 2.2045586543 | 0.1195125530 | 0.0993713219 |
| C | 6.0 | 1.1411103108 | 2.7081597187 | 0.3035376923 |
| O | 8.0 | 2.2851504531 | 2.3121770481 | 0.2169832255 |
| C | 6.0 | 0.7347130684 | 4.1519572483 | 0.4506460169 |
| H | 1.0 | 1.6121932032 | 4.7803472369 | 0.2952210378 |
| H | 1.0 | -0.0461034017 | 4.4221682472 | -0.2682189536 |
| H | 1.0 | 0.3415152617 | 4.3264311832 | 1.4595086730 |
| H | 1.0 | -1.9978810922 | 0.3601636353 | 0.2400687669 |

**g) 5fC-H+**

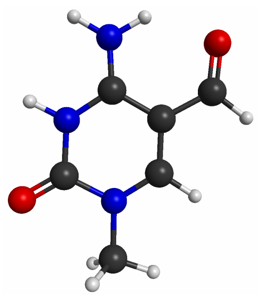

*E(LUMO) = -2.465*

| ATOM | CHARGE | X | Y | Z |
|---|---|---|---|---|
| C | 6.0 | -0.1671409289 | 0.5291291589 | 0.0771499245 |
| C | 6.0 | 1.0753581136 | -0.1847274316 | -0.0172854649 |
| C | 6.0 | 1.0176896197 | -1.5613727733 | -0.0534032386 |

| | | | | |
|---|---|---|---|---|
| N | 7.0 | -0.1312597189 | -2.2577348512 | -0.0030332535 |
| C | 6.0 | -1.3755218733 | -1.6103897814 | 0.0951226705 |
| N | 7.0 | -1.3031245994 | -0.2161990303 | 0.1264330785 |
| H | 1.0 | 1.9270148885 | -2.1491241210 | -0.1271230357 |
| C | 6.0 | 2.3659084405 | 0.4985902184 | -0.0810041460 |
| H | 1.0 | 3.2443875493 | -0.1660540084 | -0.1532603424 |
| C | 6.0 | -0.1607662234 | -3.7324532148 | -0.0451599010 |
| H | 1.0 | 0.8617065660 | -4.0910033832 | -0.1547151962 |
| H | 1.0 | -0.7653500524 | -4.0548745760 | -0.8941813280 |
| H | 1.0 | -0.5952132190 | -4.1122169850 | 0.8812982760 |
| O | 8.0 | -2.4279478257 | -2.2082449176 | 0.1467753423 |
| N | 7.0 | -0.2277176503 | 1.8429039889 | 0.1167302753 |
| H | 1.0 | 0.6567349980 | 2.3515525304 | 0.0753904716 |
| H | 1.0 | -1.1038482622 | 2.3495453620 | 0.1756239532 |
| O | 8.0 | 2.4997823794 | 1.7182279397 | -0.0586879064 |
| H | 1.0 | -2.2085405013 | 0.2432564453 | 0.1928154208 |

**h) 5caC-H+**

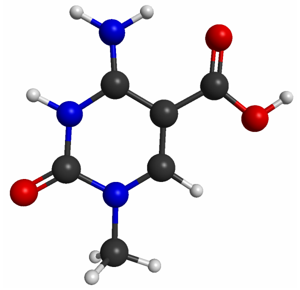

*E(LUMO) = -2.367*

| ATOM | CHARGE | X | Y | Z |
|---|---|---|---|---|
| C | 6.0 | -0.1278569711 | 0.5252773969 | 0.0757917241 |

| | | | | |
|---|---|---|---|---|
| C | 6.0 | 1.1080036761 | -0.1987776091 | -0.0033658273 |
| C | 6.0 | 1.0382154239 | -1.5740996582 | -0.0388791366 |
| N | 7.0 | -0.1185316279 | -2.2615807024 | -0.0017599974 |
| C | 6.0 | -1.3569256765 | -1.6066721956 | 0.0801102735 |
| N | 7.0 | -1.2694633697 | -0.2149339184 | 0.1112748587 |
| H | 1.0 | 1.9424241235 | -2.1672685926 | -0.1006384106 |
| C | 6.0 | 2.3987913049 | 0.5129349544 | -0.0504495252 |
| C | 6.0 | -0.1566339646 | -3.7357093919 | -0.0446266228 |
| H | 1.0 | 0.8647985457 | -4.1014642047 | -0.1390077020 |
| H | 1.0 | -0.7510083096 | -4.0546102460 | -0.9022291842 |
| H | 1.0 | -0.6072060725 | -4.1130280508 | 0.8751838314 |
| O | 8.0 | -2.4165263365 | -2.1943375609 | 0.1188814820 |
| N | 7.0 | -0.2019237274 | 1.8395447732 | 0.1152838775 |
| H | 1.0 | 0.6739186055 | 2.3613855526 | 0.0851107910 |
| H | 1.0 | -1.0858186349 | 2.3329167662 | 0.1646389283 |
| O | 8.0 | 2.5070975103 | 1.7322926727 | -0.0237100122 |
| O | 8.0 | 3.4475375473 | -0.3157050401 | -0.1261790943 |
| H | 1.0 | 4.2615185416 | 0.2253511325 | -0.1561592944 |
| H | 1.0 | -2.1706025683 | 0.2537447021 | 0.1653647504 |

**i) 5-ClC-H+**

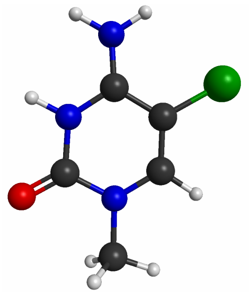

*E(LUMO) = -2.297*

| ATOM | CHARGE | X | Y | Z |
|---|---|---|---|---|

---------------------------------------------------------------

| | | | | |
|---|---|---|---|---|
| C | 6.0 | -0.1436642415 | 0.5186291491 | 0.0690864987 |
| C | 6.0 | 1.0775605891 | -0.2112723727 | -0.0288537025 |
| C | 6.0 | 1.0250858908 | -1.5769066482 | -0.0631267011 |
| N | 7.0 | -0.1416882328 | -2.2668243289 | -0.0048512212 |
| C | 6.0 | -1.3720021584 | -1.6176825023 | 0.0980311686 |
| N | 7.0 | -1.2826256904 | -0.2219835204 | 0.1241317335 |
| H | 1.0 | 1.9294320333 | -2.1686533991 | -0.1401231859 |
| C | 6.0 | -0.1735864703 | -3.7406171404 | -0.0437871764 |
| H | 1.0 | 0.8463165502 | -4.1027544342 | -0.1651582354 |
| H | 1.0 | -0.7884666416 | -4.0648018427 | -0.8850240101 |
| H | 1.0 | -0.5975515294 | -4.1200236983 | 0.8879844824 |
| O | 8.0 | -2.4365929441 | -2.1990482955 | 0.1583867286 |
| N | 7.0 | -0.2150428257 | 1.8369746240 | 0.1080885970 |
| H | 1.0 | 0.6277347800 | 2.3980427713 | 0.0664059490 |
| H | 1.0 | -1.0995026625 | 2.3281692879 | 0.1676179510 |
| CL | 17.0 | 2.5907140387 | 0.6352579628 | -0.1083630425 |
| H | 1.0 | -2.1826520756 | 0.2467016375 | 0.1953303264 |

**References:**


(1) Sas-Chen, A.; Thomas, J. M.; Matzov, D.; Taoka, M.; Nance, K. D.; Nir, R.; Bryson, K. M.; Shachar, R.; Liman, G. L. S.; Burkhart, B. W.; Gamage, S. T.; Nobe, Y.; Briney, C. A.; Levy, M. J.; Fuchs, R. T.; Robb, G. B.; Hartmann, J.; Sharma, S.; Lin, Q.; Florens, L.; Washburn, M. P.; Isobe, T.; Santangelo, T. J.; Shalev-Benami, M.; Meier, J. L.; Schwartz, S. Dynamic RNA Acetylation Revealed by Quantitative Cross-Evolutionary Mapping. *Nature* **2020**, *583* (7817), 638–643.

(2) Schmidt, M. W.; Baldridge, K. K.; Boatz, J. A.; Elbert, S. T.; Gordon, M. S.; Jensen, J. H.; Koseki, S.; Matsunaga, N.; Nguyen, K. A.; Su, S.; Windus, T. L.; Dupuis, M.; Montgomery, J. A. General Atomic and Molecular Electronic Structure System. *Journal of Computational Chemistry*. 1993, pp 1347–1363. https://doi.org/10.1002/jcc.540141112.

(3) Barca, G. M. J.; Bertoni, C.; Carrington, L.; Datta, D.; De Silva, N.; Deustua, J. E.; Fedorov, D. G.; Gour, J. R.; Gunina, A. O.; Guidez, E.; Harville, T.; Irle, S.; Ivanic, J.; Kowalski, K.; Leang, S. S.; Li, H.; Li, W.; Lutz, J. J.; Magoulas, I.; Mato, J.; Mironov, V.; Nakata, H.; Pham, B. Q.; Piecuch, P.; Poole, D.; Pruitt, S. R.; Rendell, A. P.; Roskop, L. B.; Ruedenberg, K.; Sattasathuchana, T.; Schmidt, M. W.; Shen, J.; Slipchenko, L.; Sosonkina, M.; Sundriyal, V.; Tiwari, A.; Galvez Vallejo, J. L.; Westheimer, B.; Włoch, M.; Xu, P.; Zahariev, F.; Gordon, M. S. Recent Developments in the General Atomic and Molecular Electronic Structure System. *J. Chem. Phys.* **2020**, *152* (15), 154102.

(4) Hehre, W. J.; Ditchfield, R.; Pople, J. A. Self—Consistent Molecular Orbital Methods. XII. Further Extensions of Gaussian—Type Basis Sets for Use in Molecular Orbital Studies of Organic Molecules. *J. Chem. Phys.* **1972**, *56* (5), 2257–2261.

(5) Hariharan, P. C.; Pople, J. A. The Influence of Polarization Functions on Molecular Orbital Hydrogenation Energies. *Theor. Chim. Acta* **1973**, *28* (3), 213–222.

(6) Kohn, W.; Sham, L. J. Self-Consistent Equations Including Exchange and Correlation Effects. *Phys. Rev.* **1965**, *140* (4A), A1133–A1138.

(7) Becke, A. D. Density-functional Thermochemistry. III. The Role of Exact Exchange. *The Journal of Chemical Physics*. 1993, pp 5648–5652. https://doi.org/10.1063/1.464913.

(8) Stephens, P. J.; Devlin, F. J.; Chabalowski, C. F.; Frisch, M. J. Ab Initio Calculation of Vibrational Absorption and Circular Dichroism Spectra Using Density Functional Force Fields. *J. Phys. Chem.* **1994**, *98* (45), 11623–11627.

(9) Hertwig, R. H.; Koch, W. On the Parameterization of the Local Correlation Functional. What Is Becke-3-LYP? *Chem. Phys. Lett.* **1997**, *268* (5-6), 345–351.

(10) Miertuš, S.; Scrocco, E.; Tomasi, J. Electrostatic Interaction of a Solute with a Continuum. A Direct Utilizaion of AB Initio Molecular Potentials for the Prevision of Solvent Effects. *Chem. Phys.* **1981**, *55* (1), 117–129.

(11) Li, H.; Jensen, J. H. Improving the Efficiency and Convergence of Geometry Optimization with the Polarizable Continuum Model: New Energy Gradients and Molecular Surface Tessellation. *J. Comput. Chem.* **2004**, *25* (12), 1449–1462.

(12) Tomasi, J.; Mennucci, B.; Cammi, R. Quantum Mechanical Continuum Solvation Models. *Chem. Rev.* **2005**, *105* (8), 2999–3093.

(13) Li, H. Quantum Mechanical/molecular Mechanical/continuum Style Solvation Model: Linear Response Theory, Variational Treatment, and Nuclear Gradients. *J. Chem. Phys.* **2009**, *131* (18), 184103.

(14) Bode, B. M.; Gordon, M. S. MacMolPlt: A Graphical User Interface for GAMESS. *J. Mol. Graph. Model.* **1998**, *16* (3), 133–138, 164.

(15) Thomas, J. M.; Briney, C. A.; Nance, K. D.; Lopez, J. E.; Thorpe, A. L.; Fox, S. D.; Bortolin-Cavaille, M.-L.; Sas-Chen, A.; Arango, D.; Oberdoerffer, S.; Cavaille, J.; Andresson, T.; Meier, J. L. A Chemical Signature for Cytidine Acetylation in RNA. *J. Am. Chem. Soc.* **2018**, *140* (40), 12667–12670.

(16) Sinclair, W. R.; Arango, D.; Shrimp, J. H.; Zengeya, T. T.; Thomas, J. M.; Montgomery, D. C.; Fox, S. D.; Andresson, T.; Oberdoerffer, S.; Meier, J. L. Profiling Cytidine Acetylation with Specific Affinity and Reactivity. *ACS Chem. Biol.* **2017**, *12* (12), 2922–2926.
(17) Liu, Y.; Siejka-Zielińska, P.; Velikova, G.; Bi, Y.; Yuan, F.; Tomkova, M.; Bai, C.; Chen, L.; Schuster-Böckler, B.; Song, C.-X. Bisulfite-Free Direct Detection of 5-Methylcytosine and 5-Hydroxymethylcytosine at Base Resolution. *Nat. Biotechnol.* **2019**, *37* (4), 424–429.
(18) Van Haute, L.; Dietmann, S.; Kremer, L.; Hussain, S.; Pearce, S. F.; Powell, C. A.; Rorbach, J.; Lantaff, R.; Blanco, S.; Sauer, S.; Others. Deficient Methylation and Formylation of Mt-tRNA Met Wobble Cytosine in a Patient Carrying Mutations in NSUN3. *Nat. Commun.* **2016**, *7* (1), 1–10.

(19) Busan, S.; Weeks, K. M. Accurate Detection of Chemical Modifications in RNA by Mutational Profiling (MaP) with ShapeMapper 2. *RNA* **2018**, *24* (2), 143–148.
